# AutoMorphoTrack: A modular framework for quantitative analysis of organelle morphology, tracking, and image colocalization

**DOI:** 10.1101/2025.07.19.665650

**Authors:** Armin Bayati, Jackson G. Schumacher, Xiqun Chen

## Abstract

AutoMorphoTrack is an open-source Python framework for analyzing organelle morphology, centroid tracking and colocalization in multichannel fluorescence microscopy. Steps in the analysis are connected by means of shared masks and persistent identities. Fifty synthetic sequences and eight publicly available microscopy studies were used in evaluation. A lack of sufficient vesicle coverage with the original, revised configuration motivated the development of a separate frozen particle pipeline. Six movies were used in parameter estimation and six were set aside for validation within a previously assessed study containing 284 manually tracked trajectories across all 12 movies. The mean accepted-link recall on the validated movies improved to 0.696 from 0.212, when fixed top-hat preprocessing was changed to the particle pipeline. For the 50 synthetic sequences, the mean Dice coefficient improved to 0.877 with the particle pipeline from 0.532 with the original revised configuration. External masking of mitochondria yielded mean Dice of 0.369 for seven MoDL SIM patches and 0.060 for thirteen Myofibril confocal patches. Reproducibility tests using source-pinned command-line and interactive notebook routes recovered defined outputs on named fixtures. However, further validation for other organelles and membranes is warranted. Colocalization describes fluorescent overlap, and does not establish membrane contact or functional interaction.

**Graphical Abstract:** 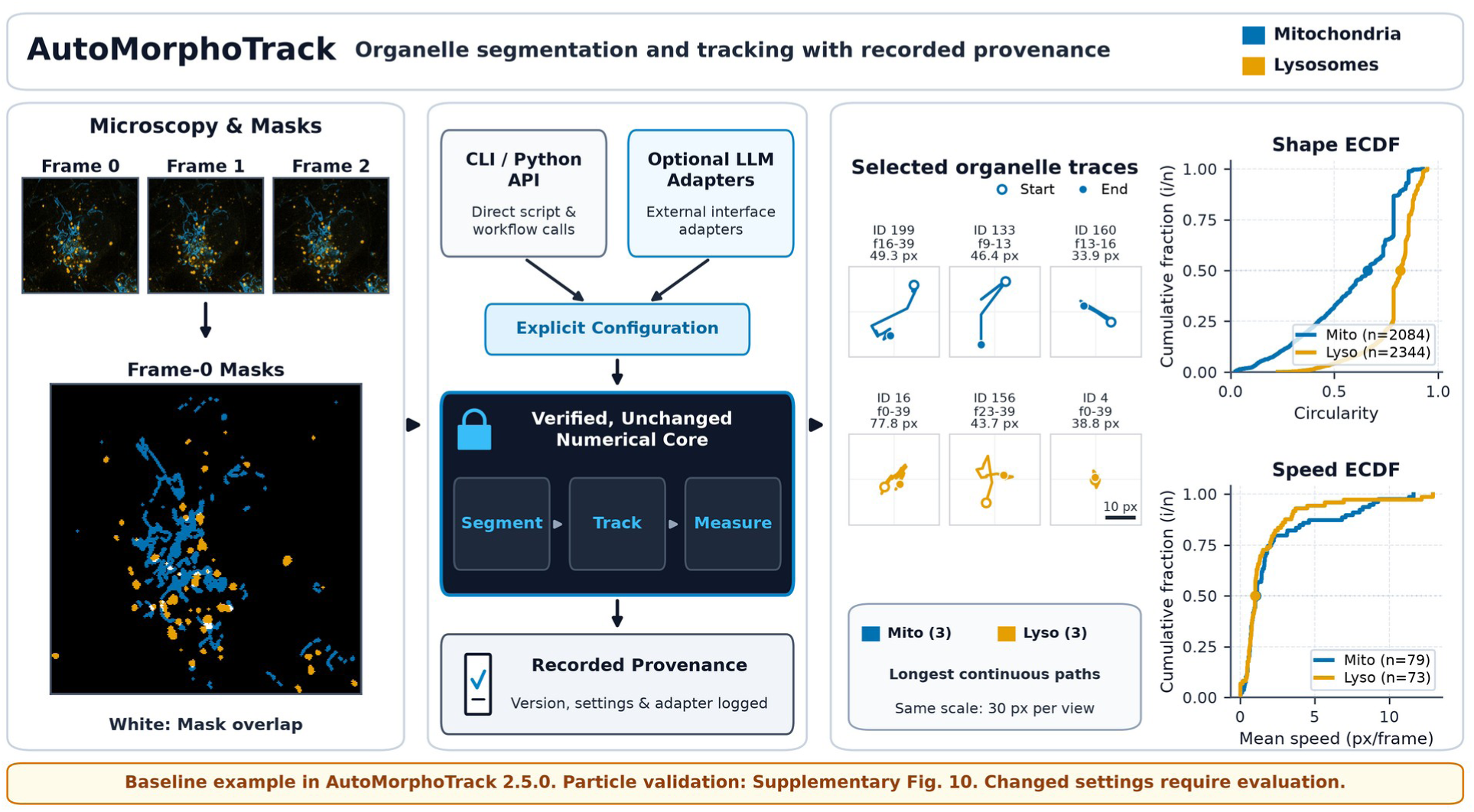

## Introduction

Quantitative live-cell imaging can show changes in the shape and movement of cellular organelles and how they are spatially organized ^1^. Dynamics and positioning of mitochondria play a role in the supply of cellular energy ^2,3^. In neuronal cells, lysosomes are involved in the transport and degradation of organelles and proteins 3. To perform an image analysis, it is necessary to distinguish the image features that are measured from the biological process that they may reflect. Many of the measurements that are needed for the analysis have already been implemented in existing open-source and commercial software. Barriers to the completion of the analysis include consistent recording of parameters, the transfer of object identity between analysis steps, and performing the analysis at an appropriate sampling level ^4–6^.

AutoMorphoTrack is a documented, Python-based workflow that performs segmentation, shape measurement, tracking, and analysis of organelle movements and colocalization. Users can perform these functions from a notebook or a Python script. The main contribution is the integration of existing analysis components using masks, object identities and recorded configuration settings. Measured parameters and quantitative results are recorded and support the reanalysis of the data and comparison to other workflows.

Users can change threshold sensitivity, minimum object size, channel assignments, and tracking distance via a configuration object. Segmentation overlays and object tables are available for visual inspection. Availability of segmentation overlays and object tables does not guarantee segmentation accuracy. The revised software captures configuration with each run to allow users to examine choices made against measured results.

Measurements saved during analysis are available for comparison between different samples, and can be combined with other image analysis workflows. Frame and organelle-level data retain their sampling structure in exported results. Examples used in the workflow evaluations and publicly available data utilize descriptive statistics to present results; treatment effects are not evaluated.

CellProfiler offers a general modular environment for image analysis. Imaris provides multidimensional visualization integrated with segmentation and tracking. MiNA and MitoGraph focus on morphology of the mitochondrial network. Cellpose and StarDist are learned instance segmentation methods with different training data and shape assumptions. AutoMorphoTrack makes use of established image-analysis methods within a smaller Python workflow. A comparison of the scope and configuration against the alternative methods is provided; a feature comparison is not a proof of superior accuracy (7-12; Supplementary Table 1). The measured accuracy comparison is between the legacy and revised AutoMorphoTrack implementations. Head-to-head accuracy against the named tools was not tested.

## Methods

### Software Architecture and Workflow Overview

AutoMorphoTrack 2.5.0 requires Python 3.11 or later and has two explicitly versioned analysis routes. The automorphotrack.validated namespace conserves the 18 modules of core 1.1.0rc2. The automorphotrack.refined namespace includes five additional modules that provide particle detection and association, described below, and make use of the unchanged baseline measurement modules. NumPy, pandas, SciPy, scikit-image, and tifffile are the main dependencies. The two implementations are distinguishable by their source, and examples show which route produced a given result.

Both the notebook and the Python interfaces use the same analysis functions. AutoMorphoTrack does not implement a chatbot to respond to natural-language prompts, interpret natural-language commands to initiate analysis, or ensure the reproducibility of automatically generated code. Reproducibility depends on saved code, input data, configuration, dependencies, and output.

### Input Data and Preprocessing

The baseline workflow analyzes two selected fluorescence channels of a 2D time series. The particle route accepts one or more selected channels and performs colocalization only for exactly two channels. Both the baseline and particle routes require channel and axis assignments. Additional organelles and imaging conditions require separate segmentation checks and validation. Support for multichannel input does not guarantee accuracy for all fluorescent structures and imaging modalities.

The loader validates the array axes and the selected channels before analysis. The loader does not automatically make the assumption that a non-singleton spatial dimension is time. Intensity values are retained for further quantitative calculations and are subject to any explicitly configured background correction. Display scaling does not impact measurement and segmentation. Interpolation or display upscaling does not provide more spatial information or establish improved segmentation accuracy.

### Adaptability

Segmentation parameters include threshold method, threshold factor, minimum object size, and morphological cleanup. The same masks are used for counting, shape measurements, tracking and mask-based colocalization. Parameter sensitivity is evaluated separately from the illustrative neuronal analysis. Bleaching, uneven background, crowding, saturation and touching objects can affect the masks and are valid failure modes.

Objects are labeled in each frame and tracked across frames. Each exported object or track-observation record retains sample, frame, and identity information; the organelle type is identified by the filename. A consistent file structure facilitates comparison of results, but does not eliminate variations in image quality, labeling, and/or design of the experiment.

**Figure 1.**
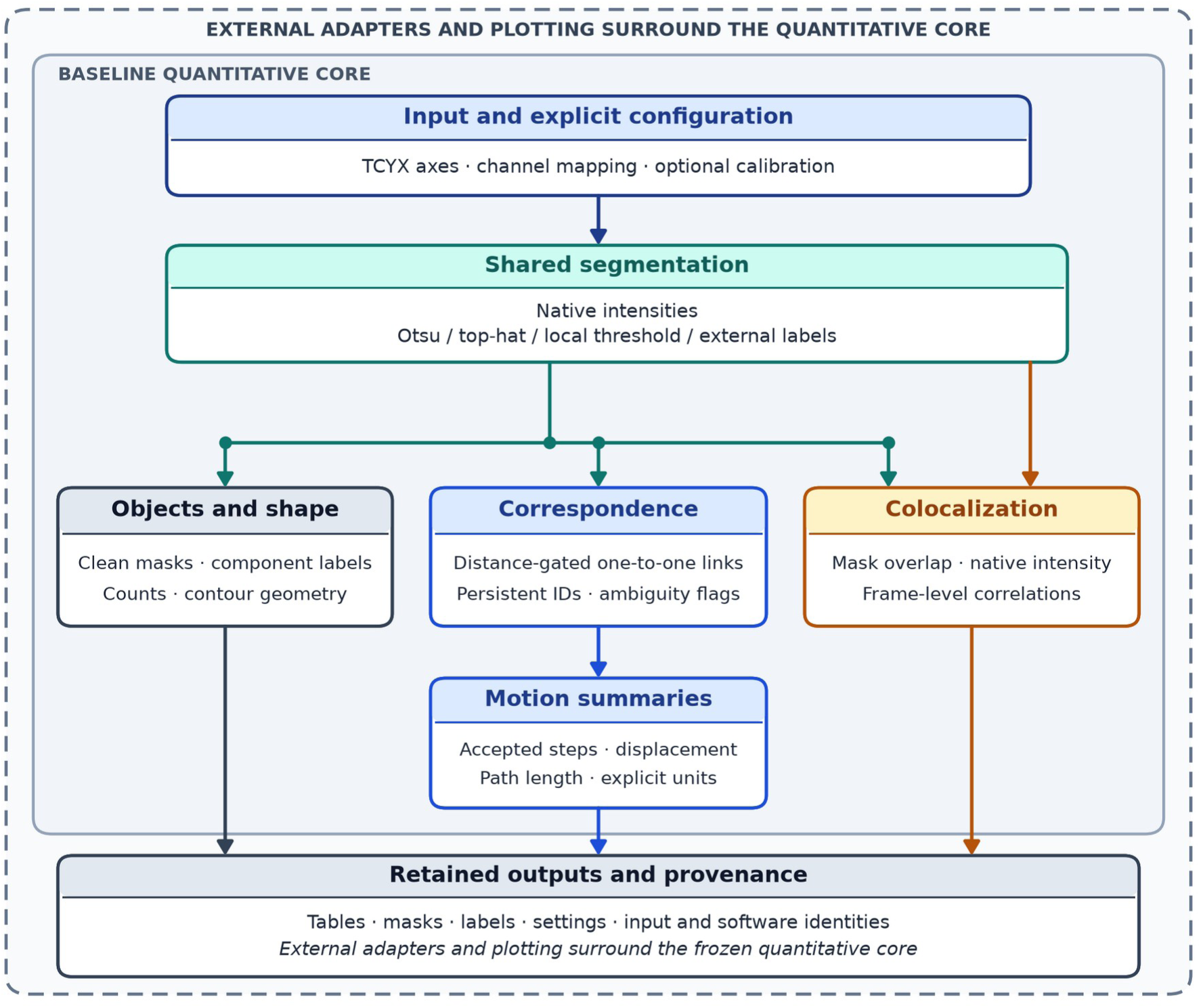
Baseline workflow in AutoMorphoTrack 2.5.0. The flowchart shows the baseline operations retained in AutoMorphoTrack 2.5.0 under automorphotrack.validated. Defined axes, channel assignments, and optional spatial and temporal calibration define the input. Native intensities are used for segmentation and intensity measurements and the same cleaned masks and component labels are used for shape measurements and mask-based colocalization, as well as for object counting and centroid linking. The assignment of one-to-one, distance-gated links produces persistent track identities and flags ambiguous links. Only accepted steps are included in the default motion summaries. The connection between inputs, settings, and measurements is preserved by tables, masks, labels and manifests. Plotting and external adapters are outside the core, quantitative functions. The separate alignment helper produces diagnostic displays and does not resample inputs used for primary measurements. The data flow described in this diagram represents functions that have been implemented. It is not a demonstration of accuracy for new data.

### Cell Culturing and Preparation

Human induced pluripotent stem cell (iPSC)–derived Neural Progenitor Cells (NPCs) were obtained from STEMCELL Technologies (Catalog #200-0620) and expanded in Neural Progenitor Medium (STEMCELL, Catalog #05833) according to the manufacturer’s instructions. After two passages, NPCs underwent midbrain differentiation using the Midbrain Neuron Differentiation Kit (STEMCELL, Catalog #100-0038) supplemented with 200 ng/mL Human Recombinant Sonic Hedgehog (Shh; STEMCELL, Catalog #78065) for 7 days. Dopaminergic NPCs were then plated on poly-D-lysine and Matrigel–coated glass-bottom 6-, 24-, or 96-well plates (Thermo Scientific, Nunc) for Western blotting, fluorescence microscopy, and quantitative imaging, respectively.

The NPCs were purchased as characterized cells. The available records do not describe an independent method of authentication or mycoplasma testing.

Differentiated NPCs were matured into neurons using the Midbrain Neuron Maturation Kit (STEMCELL, Catalog #100-0041) for two weeks. Following maturation, neurons were maintained in BrainPhys Neuronal Medium supplemented with the SM1 Kit (STEMCELL, Catalog #05792). For each biological replicate, neurons derived from independent differentiation events were plated on glass coverslips (Thomas Scientific, Catalog #1217N79) pre-coated with Matrigel (Corning, Catalog #356234) in 12- or 24-well plates. New NPC stocks were thawed and differentiated independently for subsequent biological replicates.

Fluorescence imaging was performed using a Nikon C2 confocal microscope with NIS-Elements software. Dopaminergic neurons were plated on MatTek P35G-1.5-14-C glass-bottom dishes and matured for 2 weeks. Just prior to imaging, neurons were stained with MitoTracker Orange (1:2,000 dilution; 30 minutes in BrainPhys medium; Invitrogen M7510) and LysoTracker Green (1:2,000 dilution; 30 minutes in BrainPhys medium; Invitrogen L7526). Live imaging was performed using a stage-top chamber set at 37°C and 5% CO₂. The source recording contains 40 two-channel uint16 frames. Its metadata describe a Plan Fluor 40× oil objective with NA 1.3. The reported 5 s interval conflicts with the 7.497 s interval reported in the TIFF metadata, thus, the revised motion analysis was performed using frame units. The ten examples of regions were reconstructed from the native intensity recording using the coordinates of the display crops. Channel assignments and crop coordinates are included in the input. Source hashes and explicit TCYX axes are also included. The original display crops are retained for comparison.

Steps 1 to 8 and Tables 1 to 3 describe the baseline route retained in AutoMorphoTrack 2.5.0 as automorphotrack.validated. The baseline quantitative results are shown in Figures 2, 3, and 5-17 and Supplementary Figures 1-9. The particle route, automorphotrack.refined, employs detector and association rules described in the particle refinement and reuses the baseline shape, motion and colocalization measurements. Evaluation of this route and its rules is reported in Supplementary Figure 10 and Table 9. The two routes have different configurations and output files.

**Figure 2.**
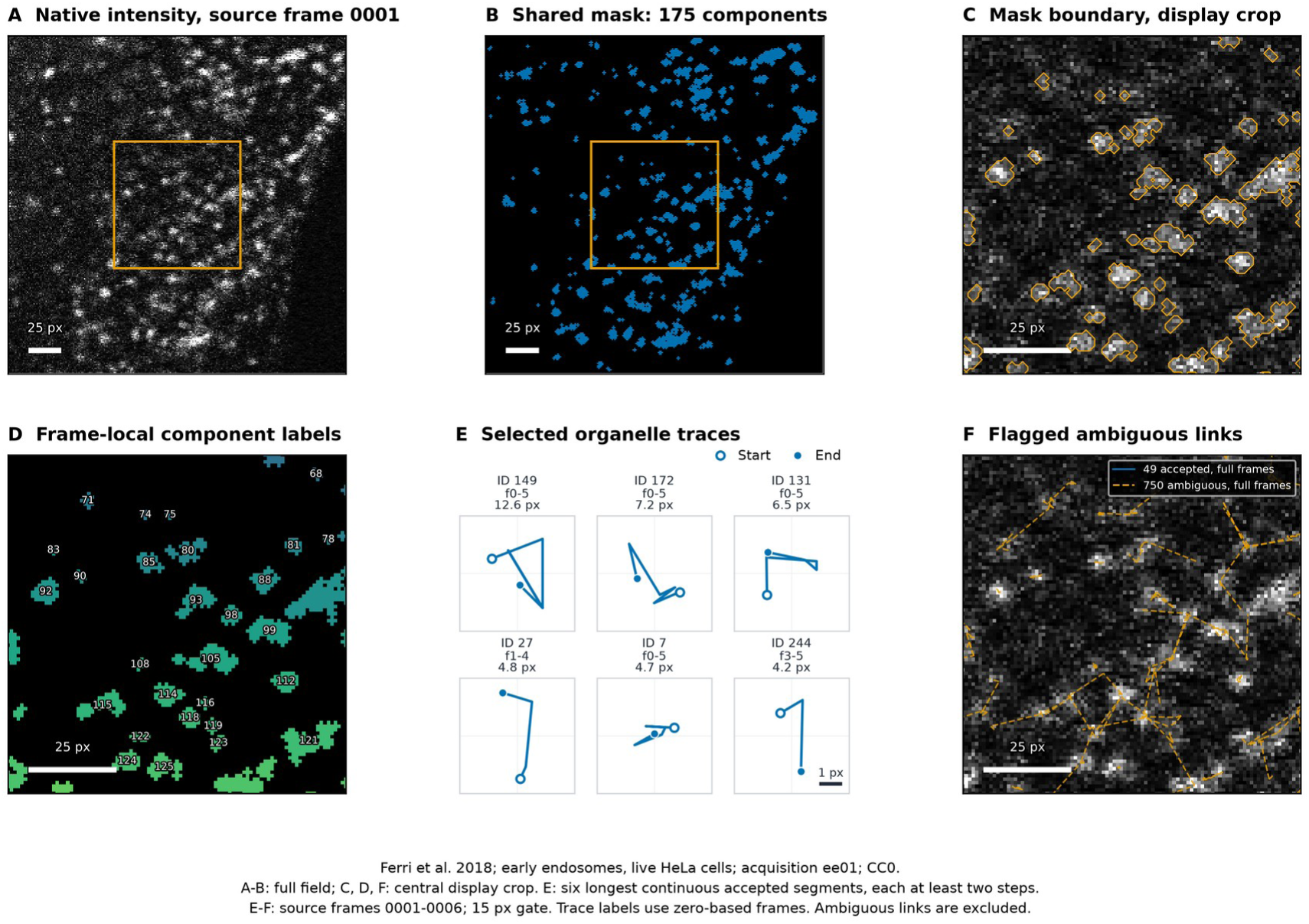
Public confocal image workflow. Public early-endosome confocal images from Ferri et al. (2018), acquisition ee01, demonstrate the revision functions on native uint16 data. A shows source frame ee01_0001.tif. B shows the foreground mask associated with ee01_0001.tif and contains 175 full frame components. Orange boxes show the central display crop (80-175 rows and 80-175 columns; zero-based indices) shown in C, D, and F. C shows the foreground mask boundaries and D labels connected components, with labels local to that frame and omitted if text would cross the edge of the display. E displays the six continuous accepted trace segments with the longest path length in source frames 0001 to 0006, each with at least two accepted steps. Traces are centered in 5-pixel-wide views at the same scale. Rotation and smoothing have not been applied. Markers identify the beginning and end of each trace; labels provide track ID, zero-based frame range, and path length. Ambiguous links and gaps end a trace segment. These examples have been chosen to illustrate movement. F shows accepted links in blue and flagged ambiguous assignments in orange dashed lines over source frame 0001. Across the full six-frame fields there are 49 accepted links and 750 flagged ambiguous assigned links; crop boundaries clip their display. The default segmentation uses Otsu factor 0.8, a one-pixel opening disk, removal of components under three pixels, border clearing and eight-connected component labeling. The link gate is set to 15 pixels. These examples show the execution and uncertainty flags. No independent masks or trajectories are provided. Images are displayed using the same frame-0001 first-to-99.5th percentile limits and do not change measurement intensity. Image scale bars are 25 pixels; trace views in E are 5 pixels wide. Data DOI: 10.6084/m9.figshare.6715550.v1; paper DOI: 10.1038/sdata.2018.191; CC0.

**Table 1.** Functional Steps in the AutoMorphoTrack Workflow.

| **Step** | **Function** | **Parameters** | **Measured output** |
|---|---|---|---|
| 1 | Shared segmentation | method, threshold_factor, min_size_px | Labeled image components |
| 2 | Object counting | Shared instance labels | Components per frame |
| 3 | Shape categories | elongated_eccentricity, punctate_eccentricity | Operational shape classes |
| 4 | Shape measurements | Shared labels, pixel_size_um | Contour geometry and region properties |
| 5 | Centroid tracking | max_link_distance_px, exclude_ambiguous_links | Accepted steps and motion summaries |
| 6 | Colocalization | ROI, shared masks, background_percentile | Intensity descriptors and mask overlap |
| 7 | Frame summary | One row per frame, Spearman | Descriptive associations |
| 8 | Sample comparison | Recorded sample and replicate identities | Descriptive sample distributions |

**Table 2.**
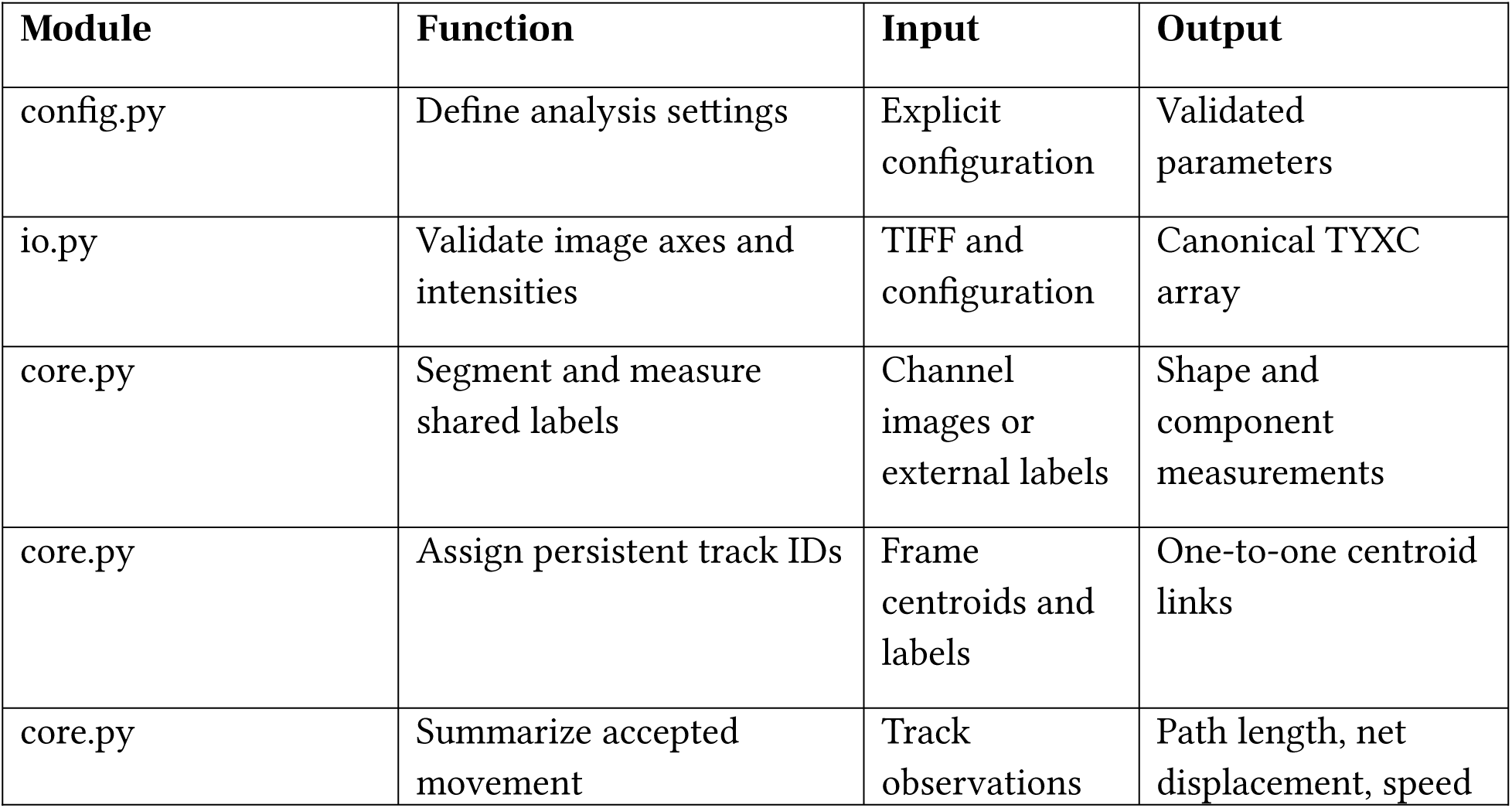
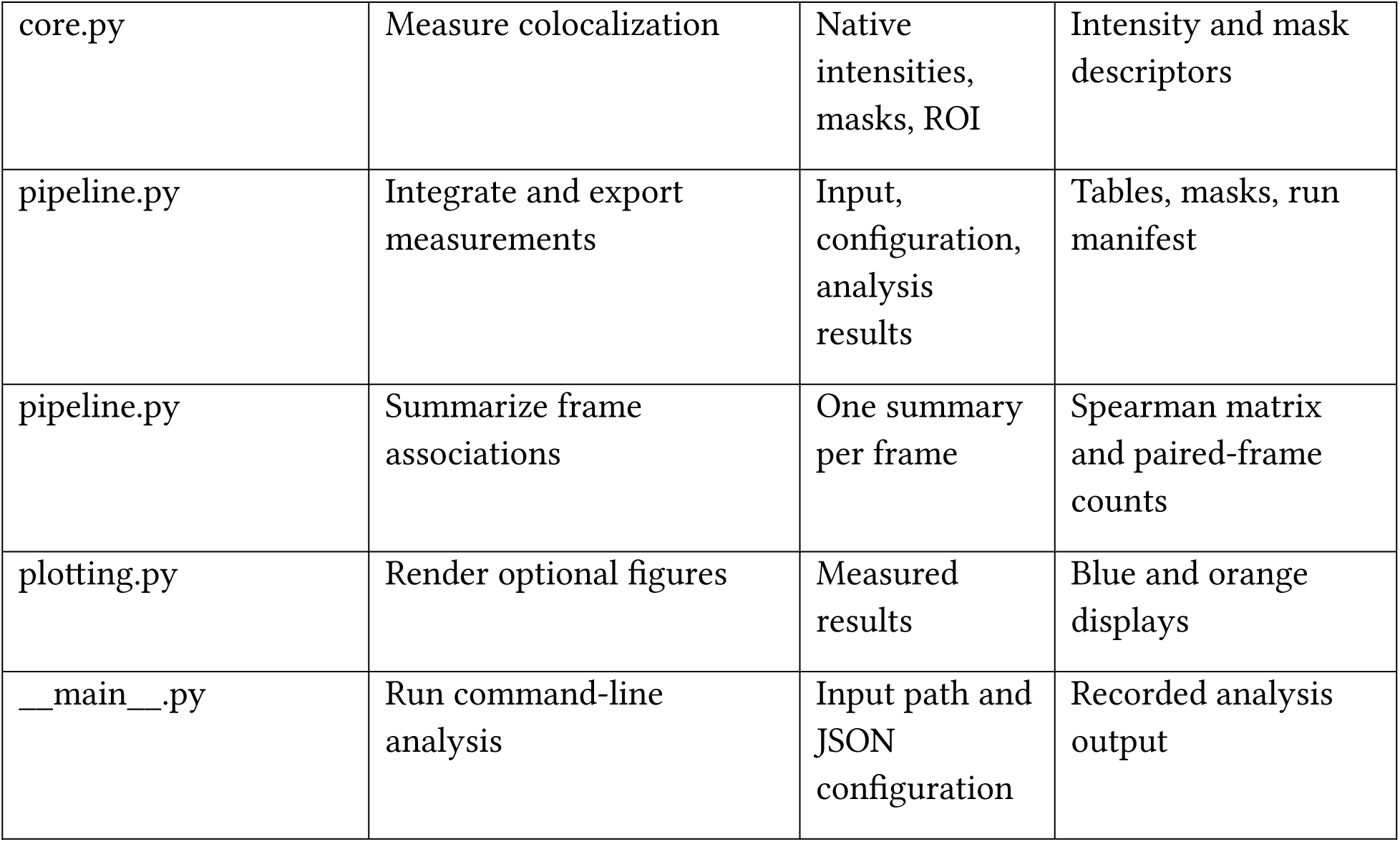
AutoMorphoTrack Package Structure.

**Table 3.**
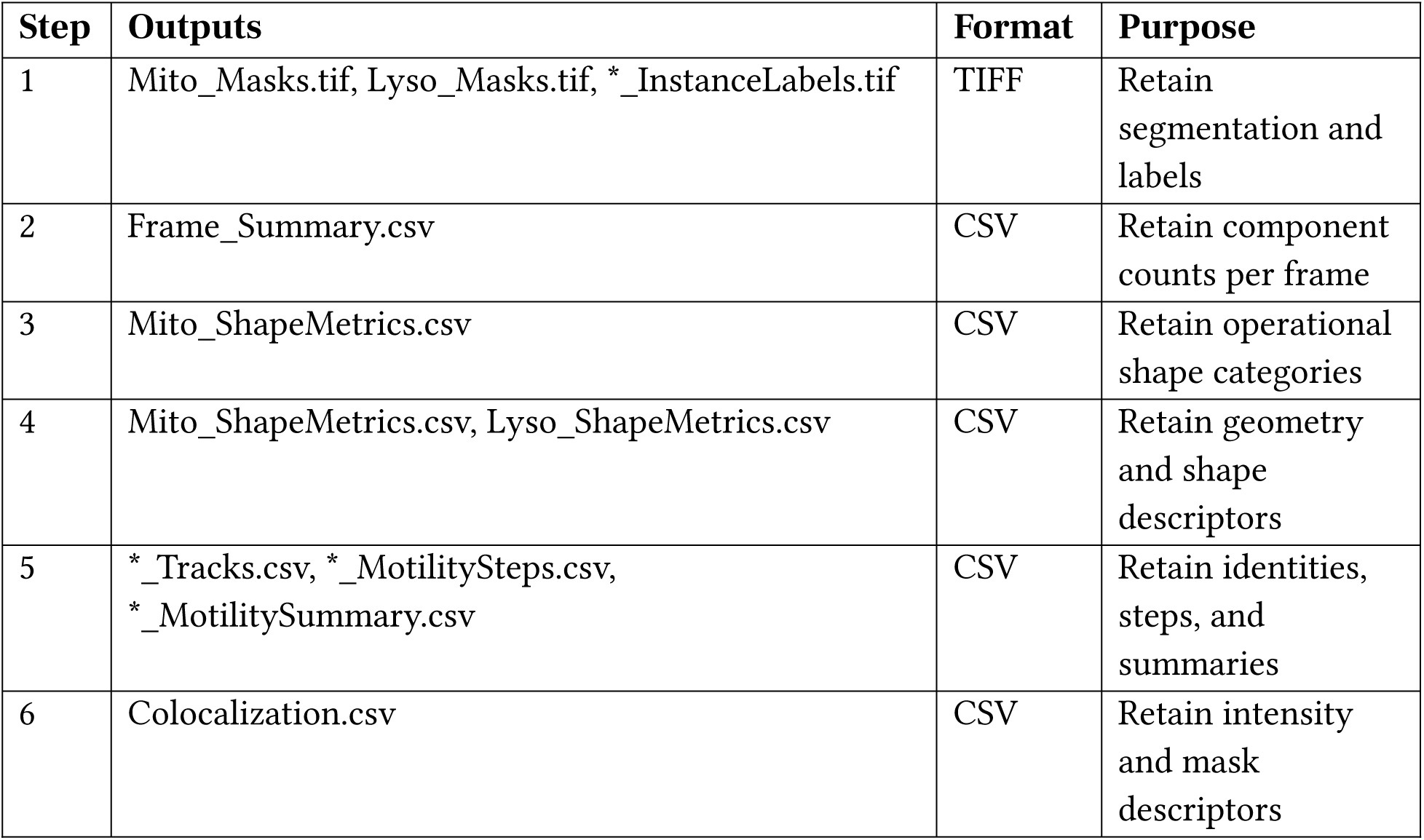
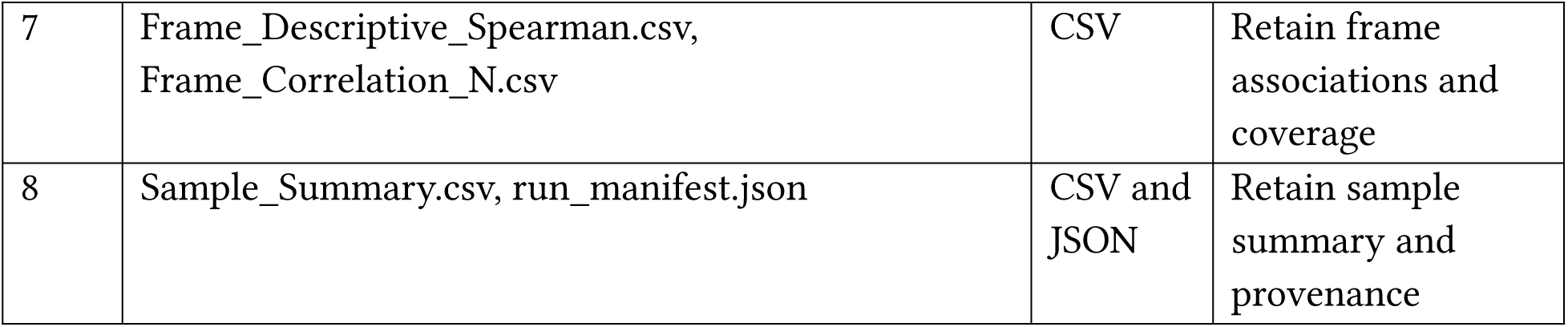
Representative Outputs Generated by AutoMorphoTrack.

| Step | Outputs | Format | Purpose |
|---|---|---|---|
| 1 | Mito_Masks.tif, Lyso_Masks.tif, *_InstanceLabels.tif | TIFF | Retain segmentation and labels |
| 2 | Frame_Summary.csv | CSV | Retain component counts per frame |
| 3 | Mito_ShapeMetrics.csv | CSV | Retain operational shape categories |
| 4 | Mito_ShapeMetrics.csv, Lyso_ShapeMetrics.csv | CSV | Retain geometry and shape descriptors |
| 5 | *_Tracks.csv, *_MotilitySteps.csv, *_MotilitySummary.csv | CSV | Retain identities, steps, and summaries |
| 6 | Colocalization.csv | CSV | Retain intensity and mask descriptors |
| 7 | Frame_Descriptive_Spearman.csv, Frame_Correlation_N.csv | CSV | Retain frame associations and coverage |
| 8 | Sample_Summary.csv, run_manifest.json | CSV and JSON | Retain sample summary and provenance |

### Step 1 – Organelle Detection

Each selected channel is segmented using thresholding and cleanup operations. The default does per-frame global Otsu thresholding multiplied by a recorded threshold factor, followed by the configured opening, small-object removal, and border handling. Optional local Gaussian thresholding and white top-hat background removal are recorded in the configuration. Users can provide their own instance-label arrays; these arrays preserve separate labeled objects that touch. Binary masks and labeled components are saved. Visual overlays are provided to aid quality control. Accuracy is assessed using reference masks, if available.

### Step 2 – Lysosome Counting

The counting module reports the number of lysosomal objects segmented in a frame. Object numbers can change due to segmentation, changes in detectability, or objects moving out of the field of view, merging or splitting. Changes in object numbers alone do not measure lysosomal degradation, formation, or turnover.

### Step 3 – Mitochondrial Morphology Classification

Shape descriptions for segmented components define mitochondrial shape categories. In the example, objects with eccentricity at least 0.8 were classified as elongated, objects with eccentricity at most 0.6 as punctate, and objects with eccentricity between these values as intermediate. No optional area gates were specified. An optional area threshold also must define a unit of measurement. For physical area thresholds, calibration of the sample distance is required. Shape categories do not indicate mitochondrial fission or fusion, or establish connectivity of the mitochondrial network.

### Step 4 – Shape Feature Extraction

Shape description and measurements include area, eccentricity, solidity, aspect ratio, orientation, and circularity of a component. Aspect ratio is the major axis length over the minor axis length of the second-moment ellipse. Orientation is the angle between the major axis of the second-moment ellipse and the row axis, in radians in the range of minus pi over two to pi over two, following scikit-image regionprops. Circularity is calculated using the equation 4πA/P², where A and P are area and perimeter of a closed contour, respectively. Pixel-count area is reported separately in square pixels. Circularity is not clipped to one. Degenerate or undefined shape measurements are reported as missing. The defined measurement replaces the inconsistent raster area and perimeter estimators in the original implementation.

### Step 5 – Organelle Tracking and Motility Analysis

Centroids are linked between consecutive frames by distance-gated one-to-one assignment using the Hungarian algorithm. Persistent track IDs are different from component labels, which are local to a frame. New tracks are started by unmatched detections. Tracks are terminated by unmatched prior objects. A link is flagged as ambiguous if either connected object has more than one candidate neighbor within the distance gate in the adjacent frame. Default motion summaries exclude these links, and so the retained trajectories generally favor locally isolated objects. The tracker does not make inferences on mitochondrial fission, fusion, or organelle lineage. The path length is the sum of accepted step distances in a track. The net displacement of a track is the distance between the start and end points of the track, reported only when all links in the observed trajectory are accepted. The mean speed of a track is the accepted path length divided by the corresponding accepted duration. The tracker reports distances in pixels and speeds in pixels per frame without confirmed spatial and temporal calibration. Video playback rate has no effect on the acquisition interval.

### Step 6 – Colocalization Analysis

Colocalization outputs are reported based on either pixel intensity or binary masks. Jaccard overlap is reported as the intersection of the two binary masks divided by the union of the two binary masks. Mask-conditioned intensity fractions are reported as the mitochondrial or lysosomal intensity within the intersection of the masks divided by the total intensity of that organelle within the mask. These values are reported along with the definitions of the masks and thresholds used. Pearson correlation is calculated from channel intensities within the shared analysis region. Intensity cosine overlap is the dot product of the two channel-intensity vectors in the shared analysis region, divided by the product of their Euclidean norms. If either norm is zero, the measure is undefined. If background_percentile is set, the specified percentile is subtracted from each channel, and any negative values are set to zero, prior to calculating intensity-based colocalization metrics. Missing denominators produce undefined fractions; constant-intensity channels make Pearson correlation undefined. Optical overlap depends on resolution, registration, background, and spectral crosstalk; it does not establish membrane contact or functional interaction.

### Step 7 – Integrated Correlation Summary

Measurements are grouped by sample, frame, and organelle type before merging. Identifiers are excluded from the correlation matrix. Spearman rank correlations indicate frame-level measurements are associated. NaN values occur with constant columns and insufficient paired measurements. Interpreting these matrices requires consideration of temporal dependence and shared derivation of the metrics. These descriptive matrices do not account for biological replication. They also do not test that a particular mitochondrial shape causes an effect on motility or overlap.

### Step 8 – Comparative Analysis (Optional External Script)

The example analyzed ten regions in a single time-lapse recording. Nested observations exist at the frame and organelle levels. Descriptive results do not utilize treatment-level p-values. To make inferential statements about sampling from replicates, the sampling unit and structure must be defined. Benchmarking results are described at the image or sequence level. Coverage and the evaluation metric are stated.

### Software Organization and Package Structure

The configuration module sets the input axes, channel assignments, segmentation, linkage, and calibration. The loader validates the image dimensions, and the analysis functions measure objects, link objects, and compute metrics. The pipeline writes the results to tables and manifests. The measurement and plotting are separate. The original comparison script shows an example of plotting results and does not provide the statistical tests or effect size measurements described in the original manuscript.

### Summary of Outputs, Functions, and Package Design

Tables 1 to 3 show the various stages of the revised processing, software components, and exported measurements. Saved masks, instance labels, track links and run manifests allow measurements to be traced through the analysis.

### Execution and reproducibility

The revised workflow sets an explicit configuration and saves the input identity, channel mapping, axes, and processing parameters, and dependency versions with each analysis. Object and track tables retain object and track identifiers so that measurements can be traced to the masks and links that produced them.

There are language models that can help users write or edit code, but they are optional tools outside the AutoMorphoTrack software. The commands generated by the models require review and testing by the users. The manuscript makes no claim that requests in natural language would reproduce an analysis or provide an implemented software interface.

The original software and outputs are maintained separately from the revised version because changes to tracking, geometry of shapes, and definition of metrics yield different numerical results. The distributed configuration, analysis code and the identified input data are required to reproduce the results.

### Public image workflow and quality control

A public confocal sequence shows shared segmentation, local component labels, and accepted versus ambiguous centroid assignments. A quality control script estimates a translation using phase correlation and resamples the moving image for visual inspection on common valid support. This script is outside the frozen quantitative package and does not alter the images used in the primary measurements. Five prespecified integer translations of one public frame serve as controls. This helper is not given the actually applied shift. These controls do not validate performance for biological deformation, interchannel registration, and native stage drift.

### External adapter integration

The example for 2.5.0 particles runs through examples/refined_verified.py in an isolated Python process. Before analysis, this example checks the installed package version, the hashes of 18 baseline and 5 particle modules against their respective source manifests, and the imported module locations. Next, it runs the particle pipeline and checks the returned particle source digest. This example does not validate the wheel checksum or enforce a fixed dependency profile. The wheel and environment archives provide separate provenance.

The particle route manifest records the input checksum, native and effective axes, configuration, source digests, Python and dependency versions, platform, output checksums, and raw and retained observation counts. Physical calibration requires an explicit source; without it, motion remains in pixels and frames. Changes to adapters, preprocessing, settings or dependencies also require fixture checks and scientific assessments. Matching source hashes does not validate these changes.

For the baseline integration test, external development applications Gemini and Codex were used (20, 21). The open Gemini application constructed an adapter from the supplied interface and fixed fixture contract. Its exact captured source was then executed locally against the installed wheel; the application itself could not access or execute the local files. Codex was used to execute standalone Python commands, Jupyter notebook, regression and comparison of output. The Gemini interface showed Carbon, but its underlying model version was not independently verified. Metadata recorded Gemini version 1.141.3.1394 and OpenAI desktop version 26.1103.32239. These are application, not model versions. Supplementary Table 5 shows each route, its commands, outputs and verification. Agreement between routes tests integration and reproducibility on named fixtures. It does not guarantee an independent assessment of biological accuracy by either language model.

For the 2.5.0 route, Gemini wrote a new adapter and Codex ran the captured source against the installed wheel. The first adapter returned a file-hash mapping where the contract required a digest string. A focused change resolved that metadata error. The corrected adapter, the Codex adapter, a standalone command line, and an executed Jupyter notebook reproduced the 19 reference particle files exactly on the local host. Altered manifests and source files were rejected before analysis. The Codex guard verifies all 23 modules across both numerical namespaces. Prior to the requests, Argon High was selected in Gemini, but the interface changed its model label after the responses; the underlying model identity was not verified. These records test software integration, and numerical reproducibility, without constituting independent biological validation by either assistant.

## AI assistance for figure construction and integration

Gemini and Codex assisted with designing figures and producing plots using Python. The plotting code was executed locally, and plotted values were checked against numerical outputs. They also assisted with external adapters and integration tests, as described above. These tests assessed software integration and the output reproduction; they provided no independent biological validation.

### Controlled and public image evaluation

Controlled tests used 50 seeded two-dimensional synthetic sequences with five realizations of each of ten conditions, and eight frames per sequence. Variation of conditions included signal-to-noise ratio, background gradients, bleaching, object density and blur. Instance masks and identities provided references for pixel Dice and intersection over union, instance detection, link precision and recall, and identity switches. The ten fixed analysis configurations included the historical exported-ID baseline, revision, parameter perturbations and an oracle instance mask control. All sequence-level results are provided. Differences in default workflows reflect both implementation and parameter changes. Oracle mask controls separate segmentation errors from correspondence; they do not establish performance on biological images.

The public data protocol fixed source file selection and analysis settings prior to scoring. It included four confocal studies comprising 14 fields/acquisitions and 130 2D inputs, as well as a separate MoDL structured-illumination image mask set (13–17). Three of the ten MoDL patches were designated as development examples, and the other seven were not used for the development of AutoMorphoTrack. The source paper describes manual annotation of its segmentation dataset, but annotator histories and source-cell groupings are not available. Source model training examples are not a test set for MoDL. Color-coded images were converted to grayscale for both workflows, following the source loader. The confocal data lacked independent organelle masks and trajectories. Thus, the assessments focused on execution, numerical geometry and parameter-dependent outputs. Spatial projections, selected channels, licenses, source versions, hashes, and configuration values are provided with the benchmark.

The initial confocal panel used endolysosomal recordings from Ferri et al. (13; collection DOI: 10.6084/m9.figshare.c.4070033), MitoLandscape examples from Negri et al. (14; dataset DOI: 10.5281/zenodo.16037033), OrganelleProfiler images from Laan et al. (15; dataset DOI: 10.5281/zenodo.7875537), and mtFociCounter dataset 02 from Rey et al. (16; dataset DOI: 10.5281/zenodo.7634536). These sources provided six time-lapse acquisitions, one static mitochondrial projection, three fields consisting of five channel images, and four static mitochondrial projections, respectively. The illustrations of segmentation and controlled alignment use Ferri acquisition ee01 (early-endosome; dataset DOI: 10.6084/m9.figshare.6715550.v1). The comparison using structured illumination uses the ten MoDL demo image-mask pairs from Ding et al. (17), in repository commit 1053e473785560bc473aa64fd357379f9f0e21e2. MoDL patches 0000-0002 were development examples, and patches 0003-0009 were not used in the development of AutoMorphoTrack. The source images and analysis outputs are presented together so that segmentation behavior can be seen with the numerical analyses.

We extended the public panel with two further studies under a protocol fixed before scoring (18, 19). The Myofibril study provided the 13 published 128 by 128 pixel confocal image-mask pairs, where mitochondrial outer membranes were labeled by MOM-GFP (18). The authors manually drew the masks in Fiji. These patches came from one parent stack and were training examples for the source model and were external to the development of AutoMorphoTrack. The images and masks can be found in the author’s repository at commit 11f512a8181fb98540b6e5a89466ed85b46b4d5f. The Mito Hacker study provided all 17 original multicell TIFF fields at commit 18765f1d584db04be77f3397f730b98a51225394 (19). We used the green channel of the RGB exports, which represented mitochondria labeled by YFP. Four existing configurations (the historic baseline, revised default, fixed top-hat, and fixed local thresholding) were used in the study without training or parameter tuning. We kept patch-level Dice and intersection over union for Myofibril masks and execution status, component counts, and foreground fraction for all the input datasets. Mito Hacker provided no independent organelle masks or reference trajectories.

Biological tracking was assessed using the MSP-tracker vesicle benchmark from the Drosophila follicular epithelium, using Airyscan and spinning-disc microscopy (23, 24). We used version 1 of the deposit because benchmark files in the version 2 bundle were absent. The 12 deposited movies contained 284 manual trajectories, 5,258 localizations, and 4,959 adjacent reference links. The authors of the source article reported 391 trajectories. We scored only the deposited annotations and ignored a duplicate of the Fas3 export. JSON and XML coordinates agreed, after converting row-column to x-y order. Annotation frames were checked against each of the 12 movies. The historical baseline, revised default, a 6-pixel linking gate, and fixed top-hat configuration were set prior to scoring. Predictions were matched one-to-one to manual trajectory points within an analyst-defined 3-pixel radius. Sensitivity analyses at 2 and 5 pixels were also performed. Unmatched predictions were not verified since annotations did not exhaust all visible entities. Conditional link precision was used to assess link predictions whose two endpoints were manually annotated. This dataset is for secretory vesicles and does not provide mitochondrial or lysosomal reference identities.

Poor vesicle coverage resulted in a different version of the particle pipeline in AutoMorphoTrack 2.5.0. We skipped MSP04 to MSP06 and MSP10 to MSP12 during parameter selection and used the rest of the 12 movies for development. The 12 movies had been previously assessed with the older fixed configurations, so these split tests assess performance within the reserved movies of the same study, rather than performance in an unseen study. Nine detector settings were evaluated on every fifth development frame. Later, 16 detector and association combinations were assessed on all development frames and 9 synthetic movies with complete point references. Candidate selection required a mean synthetic detection precision of at least 0.90, a mean accepted-link precision of at least 0.95, no more than one retained noise detection per frame and a combined conditional biological accepted-link precision of at least 0.98. Of the settings which met the candidate selection criteria, we selected the settings with the highest mean development accepted-link recall and set the configuration and source hashes before assessing the reserved movies.

The selected detector subtracts a Gaussian background with a 6-pixel standard deviation and applies a scale-normalized Laplacian of Gaussian at 1.5 pixels. Peaks must exceed three times the median-absolute-deviation estimate of response variation and be separated by at least three pixels. Positive foreground is divided into instances using a seeded watershed. Each support mask is constrained to a radius of 4.5 pixels, at least 20% of the peak foreground signal, and at least three pixels. Intensity-weighted centroids of the support masks define positions. These support masks describe locations where the signal is supported. In the current implementation of this algorithm, boundaries of the support masks have not been validated as organelle membranes. Association minimizes squared, distance-normalized assignment cost, with unmatched alternatives and a 6-pixel gate. A link is only accepted if its distance is more than 0.5 pixels better than alternatives, in both assignment directions. Tracks are retained if they contain at least three observations. Tracks with fewer than three observations are excluded from motion summaries and retained masks. Raw detections are not excluded. The selected configuration does not use motion prediction, and gap closing is turned off. Ambiguous links are kept, and are excluded from speed estimates.

The particle route saves channel-indexed raw and retained label TIFFs, raw and retained track tables, shape measurements, motion steps and summaries, per-frame motion summaries, and detection thresholds. Track and shape rows are linked by frame and local label. With two selected channels, colocalization.csv records full-field measurements of the retained masks and native intensities. The output filenames are different from the baseline outputs in Table 3.

The metric for fragmentation was defined as the number of additional correctly localized contiguous segments within each continuously annotated reference trajectory. A segment was broken by (i) a missing localization, or (ii) a change in the predicted identity. Accepted-link segments were broken at an excluded ambiguous link. Gaps in the reference annotations were not considered to be errors of the algorithm. We reported fragmentation, link recall, and localization recall, and completely missed trajectories, because an undetected trajectory contains no measurable fragments. The original synthetic fixtures and settings were used, and both original link-recall measurements were reproduced across all 500 sequence-configuration evaluations.

Runtime for the baseline core, retained unchanged in 2.5.0, was measured using three fresh Python processes per rendering setting on an Apple M5 Pro with 48 GiB of memory. Each process imported the package and loaded the two-channel 40-frame Sample8 stack, performed analysis and drift auditing, and exported results to CSV and TIFF, and wrote a manifest. Rendering PNG images was done separately. Wall time was measured, including interpreter startup and shutdown. Peak resident memory was read from the OS. File caches were not cleared.

## Results

We evaluated the workflow using 50 synthetic control sequences and eight publicly available microscopy studies. Six confocal studies provided a total of 160 two-dimensional inputs from 32 fields and/or acquisitions, including 13 manually annotated patches from a single parent stack. A structured-illumination study provided seven MoDL patches, which were not a part of the AutoMorphoTrack development. Another confocal study provided 12 vesicle movies containing a total of 284 manual trajectories for biological tracking evaluation. Synthetically controlled accuracy, external mask comparisons, and execution checks are illustrated in **Figures 5 to 7**. Publicly available microscopy studies illustrated segmentation and alignment diagnostics in Figures 2 and 3 respectively. Figures 8 to 17 and **S1** to S8 illustrate examples of analysis performed on the neuronal recording.

**Figure 3.**
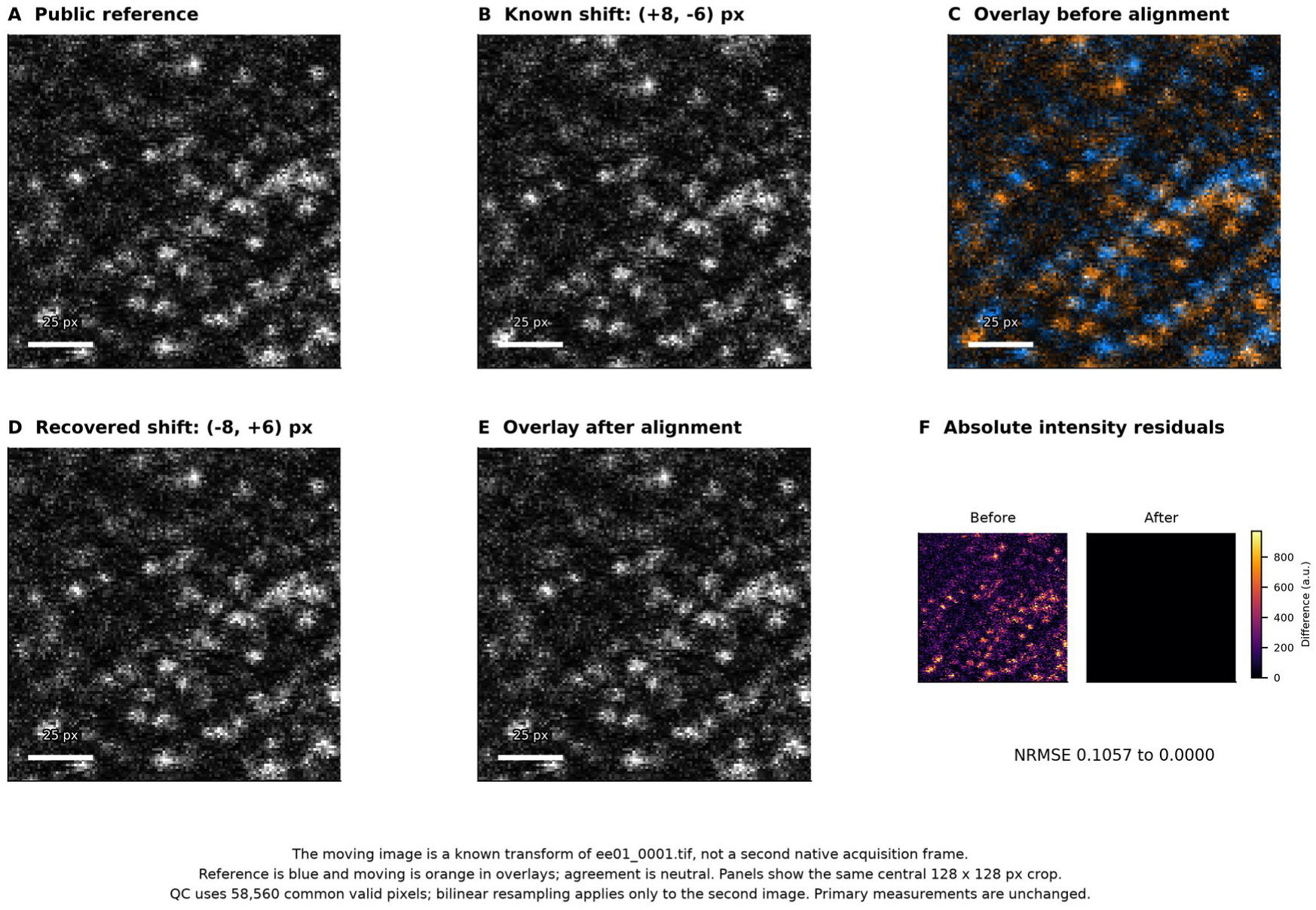
Controlled translation quality control. A controlled translation was performed on the public ee01_0001.tif frame to showcase a separate, stand-alone visual QC helper. See Image A; image B is the same as image A, but has a shift of +8 rows, -6 columns, with constant padding. Image C is the pre-alignment overlay. The estimator receives only the image arrays and does not take into account the known shift; the correction is (-8, 6) in row-column order. Image D is the resampled second image, and E is the overlay. In the overlay, agreement is represented as a neutral color, reference-only signal is in blue, and moving-only signal is in orange. F shows absolute residuals using the same intensity scale. The normalized RMSE over 58,560 valid pixels changes from 0.1057 to 0.0000. True validity excludes original padding and resampling borders, and both measurements of error use the same support. Displays show rows and columns 64-191 of the original 256x256 frame and contain a 25 pixel scale. A total of five of these prespecified integer shifts of a single image were tested; their results, along with prespecified 0.1 pixel-shift tolerance, are provided. This transform test does not prove the accuracy of the registration for biological deformation, interchannel registration or native acquisition stage drift. The helper is separate from frozen AutoMorphoTrack 1.1.0rc2, and no aligned image feeds the primary measurements. The source and license are the same as shown in Figure 2.

**Figure 4.**
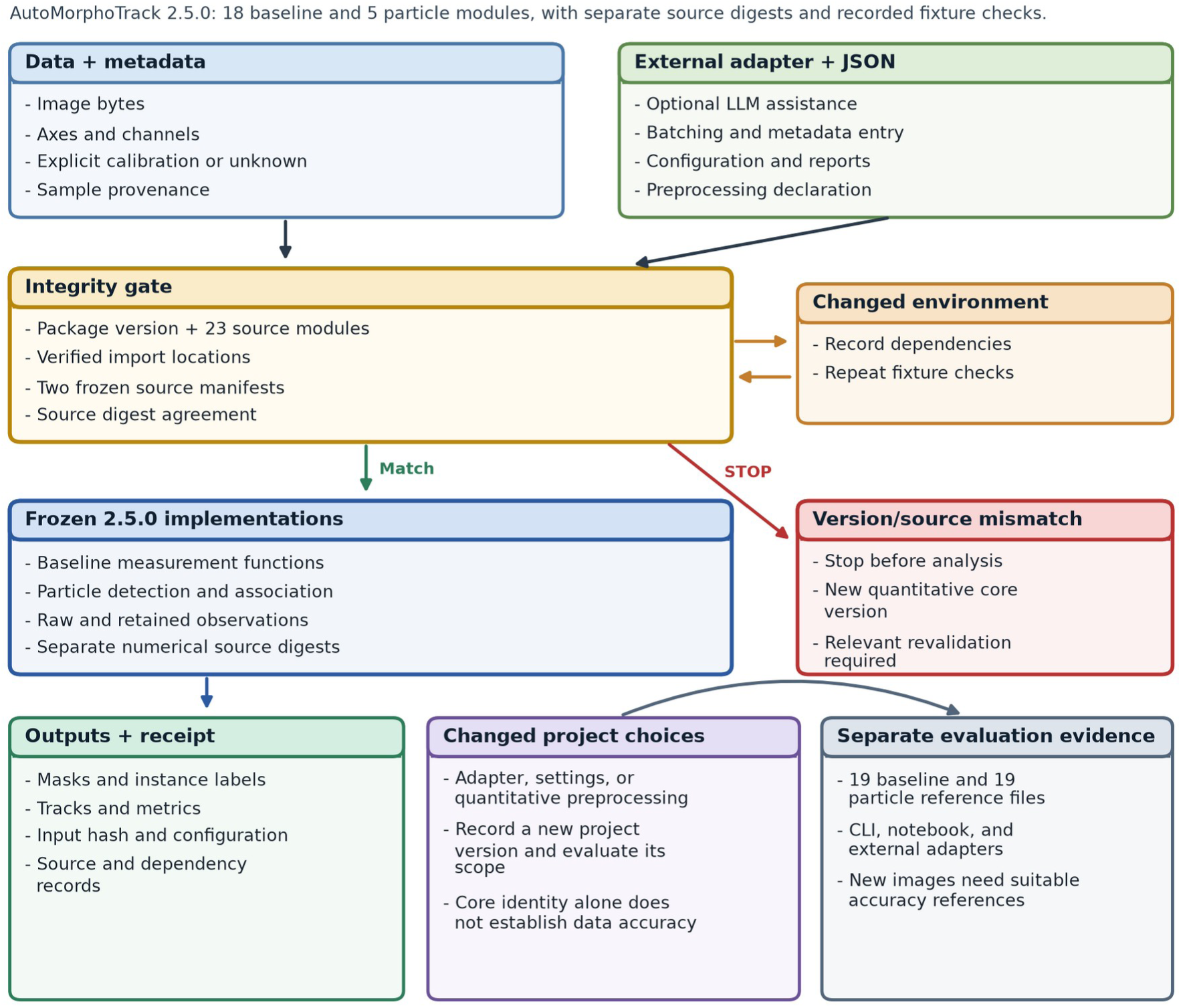
External adapter integration architecture. The input data and metadata are routed to the 2.5.0 particle pipeline via an external adapter using JSON configuration. An example helper checks the installed package version, both source manifests covering 23 modules, and import locations. A source mismatch stops the analysis. The run manifest captures input identity, settings, source and dependency versions, observation counts, and output checksums. New adapters, settings, preprocessing, and environments require new checks. The tested execution paths are supported by the supplied reference outputs; however, they do not validate output correctness for new data.

**Figure 5.**
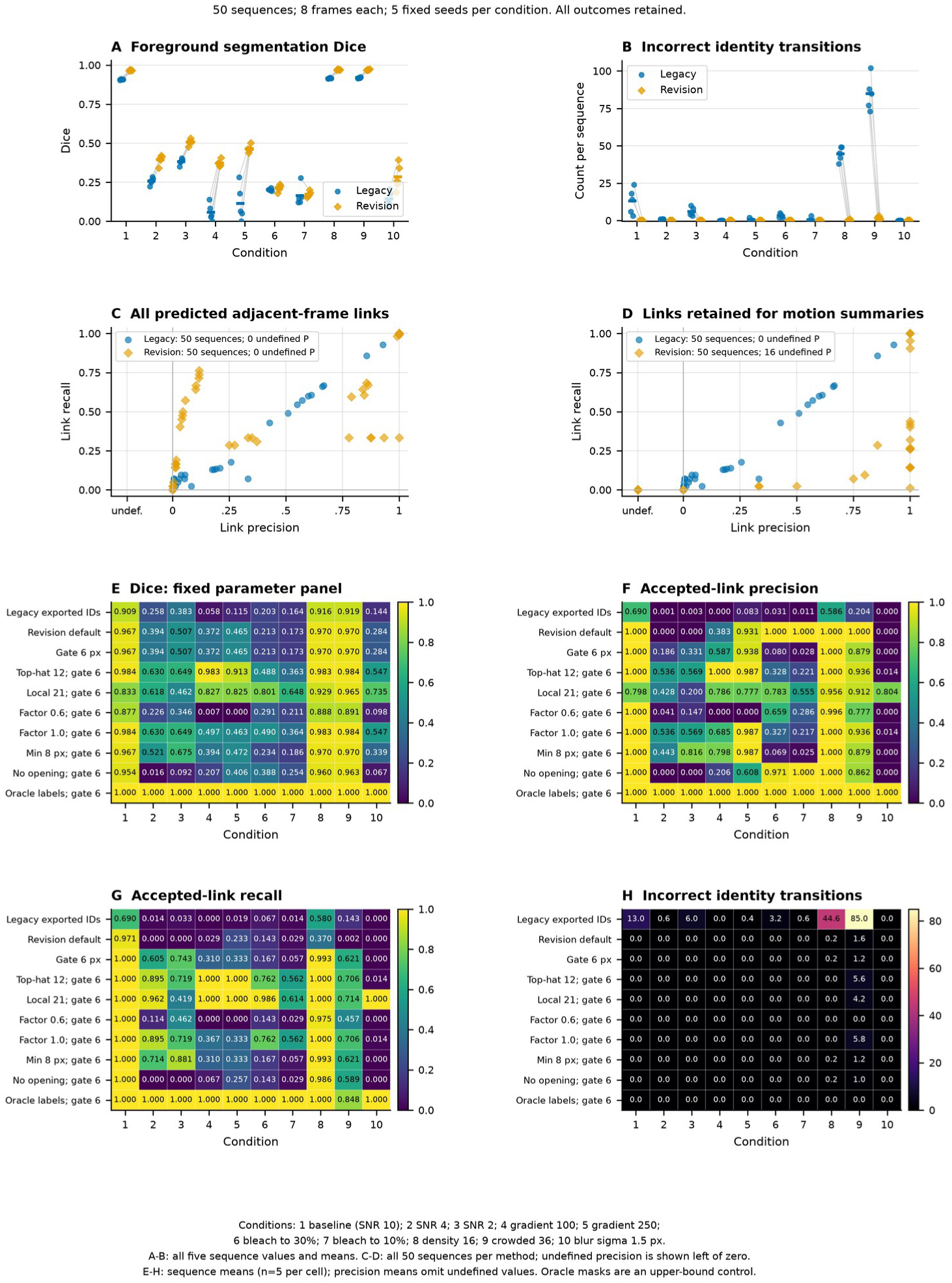
Controlled synthetic stress panel. A and B show all five seeded sequence values per condition for reconstructed legacy exported IDs and the revision default. C and D show precision and recall for 50 sequences both before and after ambiguity was removed. Undefined precision is shown to the left of zero. E through H show the ten fixed strategies/configurations for the ten conditions, with five sequences per cell. Means (averages) omit undefined precision. Finite counts are shown in the source tables. In the foreground, Dice compares predicted binary unions to what was rendered. Instance matches are one-to-one IoU >= 0.5. Correct identity links are shown between two matched detections with the same rendered identity, and precision and recall divide by the total number of linked predictions and available adjacent frame truth, respectively. Incorrect identity transitions are counted as predicted adjacent-frame links whose two matched detections have different truth identities, and must be read in conjunction with precision and recall, as they are reduced by the presence of either rejected or absent links. Oracle instance masks are a tracking upper-bound control and not a segmentation validation. The shapes designed by the analyst do not prove biological validity or general superiority.

**Figure 6.**
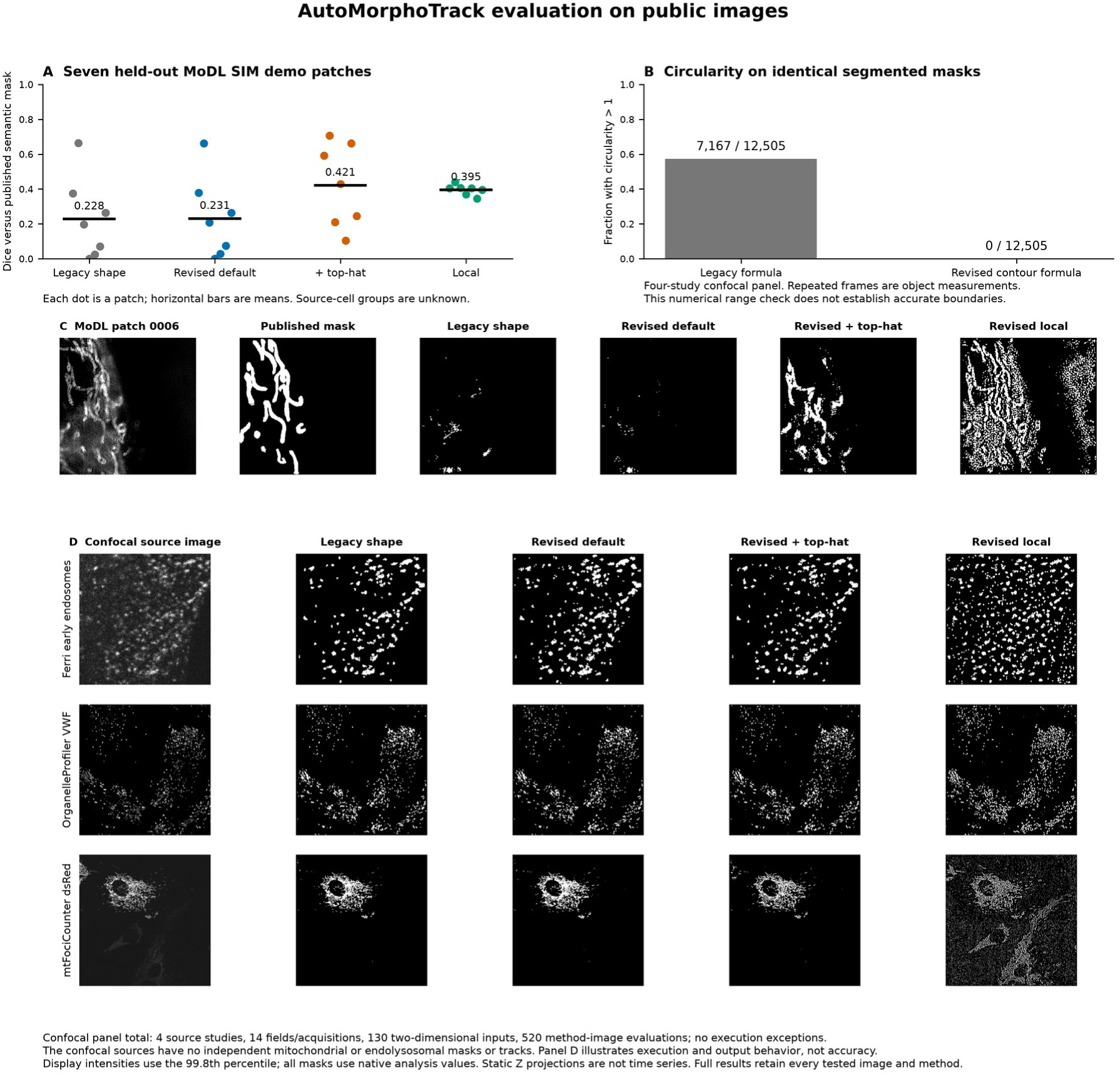
Independent public image evaluation. (A) Dice scores against published semantic masks for seven MoDL SIM demo patches held out from AMT development. Each dot represents a patch and the horizontal bars represent means. As source-cell grouping is unavailable, biological independence and absence of source-cell leakage are not established. (B) Circularity was computed using the original pixel-area/perimeter formula and the interpolated-contour formula on identical legacy segmentation masks from the complete four-study confocal panel. Using the original formula, 7,167 of 12,505 object measurements exceeded 1; using the interpolated formula, no measures exceeded 1. Repeated frame measurements are not unique organelles or biological replicates. This is a numerical consistency check, not boundary accuracy. (C) The prespecified MoDL patch 0006 shows examples of incomplete segmentation and sensitivity to background. (D) Examples include Ferri early endosomes, OrganelleProfiler VWF, and mtFociCounter mitochondrial dsRed. The complete confocal panel includes MitoLandscape, and comprises 14 fields or acquisitions, 130 2D inputs, and 520 completed method-image evaluations. Independent confocal masks or trajectory identities were not provided, thus the successful execution of the process cannot confirm segmentation or tracking accuracy. The legacy shape module uses Otsu factor 0.85 and minimum size 5 pixels. The revised default uses Otsu factor 0.8 and minimum size 3 pixels; the fixed top-hat variant adds a background filter of radius 15 pixels, and the local variant uses Gaussian local thresholding with block size 31 and factor 1. All variants use radius-1 opening and border clearing. Image displays are scaled to the 99.8th percentile; masks, however, use unscaled analysis arrays. Z projections are static. Full per-image outputs retain all tested configurations. Color-coded MoDL demo images are exports converted to grayscale using the source loader convention. The exact image-mask files are from the GPL-3.0 MoDL repository. No separate per-image license is stated. Other datasets are CC0 (Ferri) or CC BY 4.0 (OrganelleProfiler and mtFociCounter).

**Figure 7.**
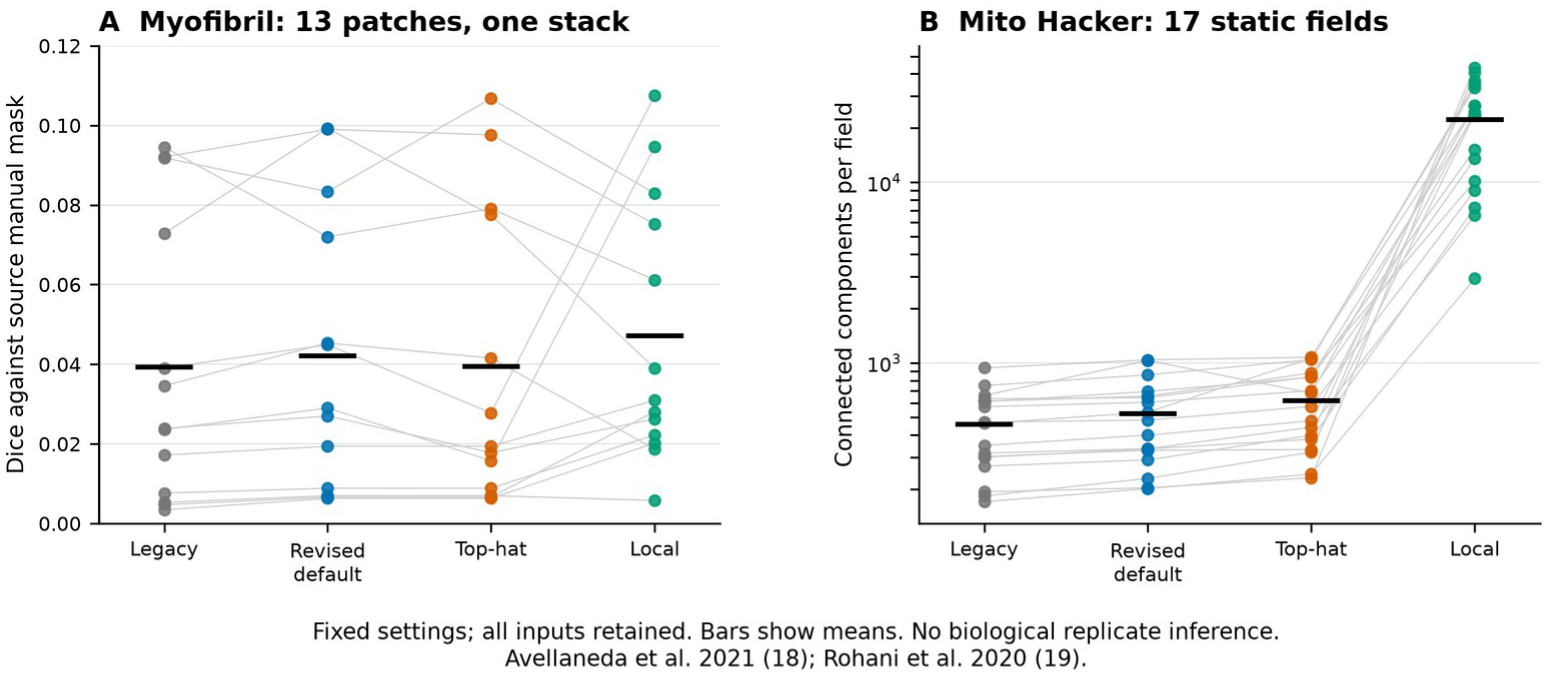
Additional confocal studies. A shows paired Dice values for all 13 manually annotated Myofibril patches from one parent stack, with horizontal bars indicating means. The four settings were fixed prior to scoring. The source patches were training examples for the source model and were external to AutoMorphoTrack development (18). B shows component counts for all 17 Mito Hacker multicell fields, analyzed through the green channel of the supplied RGB exports (19). The logarithmic vertical scale accommodates the large increase with local thresholding. There are no independent organelle masks for these fields, so component counts are descriptive. Points represent patches or fields, not independent animals or experiments. The source images are available through the cited repositories, and the exact selections, hashes, masks, and per-image measurements are retained with the analysis.

**Figure 8.**
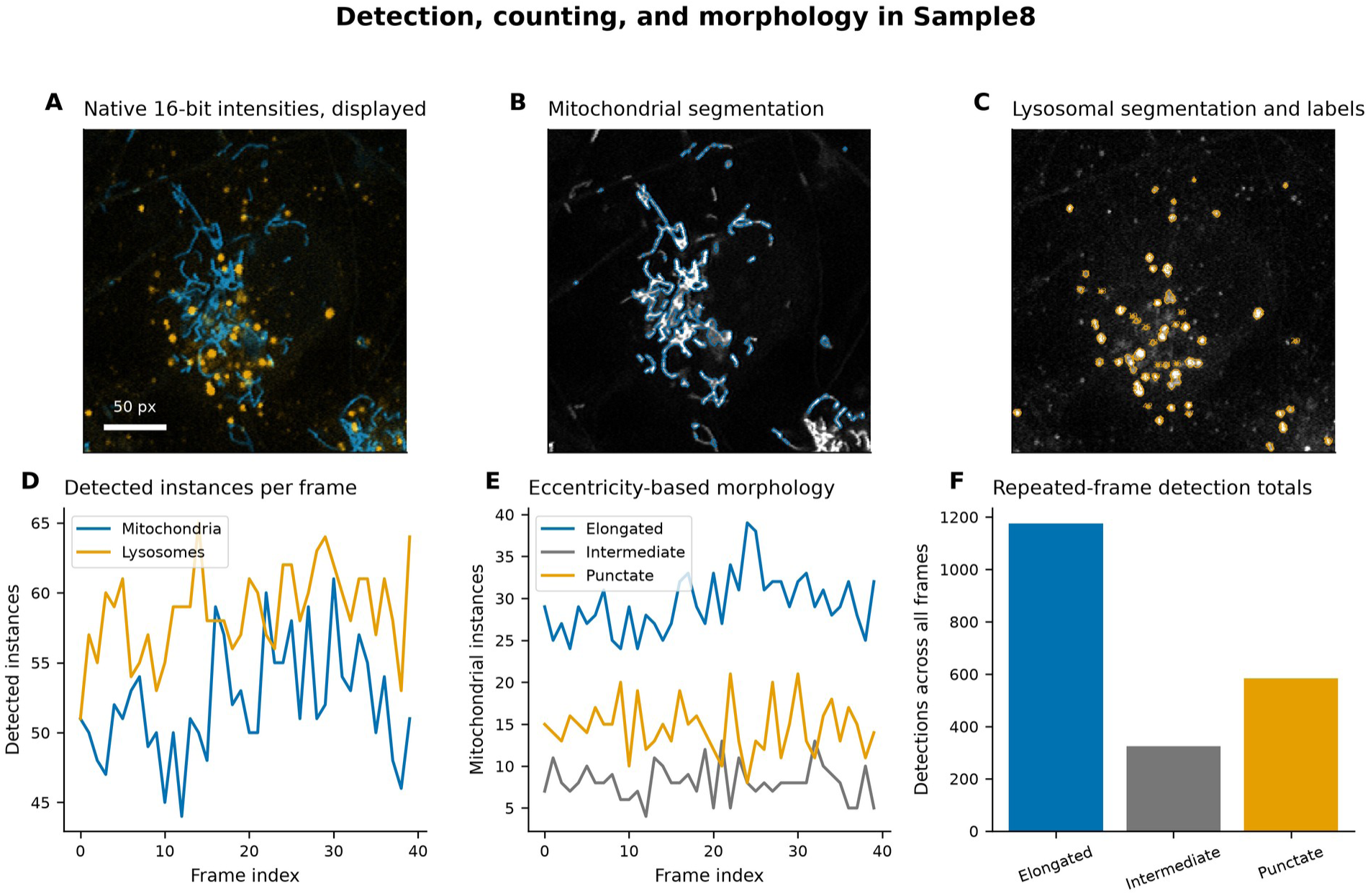
Detection counting and morphology. Sample8 contains 40 frames from one recording and encompasses 258 × 258 px. **A** through C show native intensities, segmentations of mitochondria, and segmentations of lysosomes with component labels at frame 0. The scale bar is 50 px and mitochondria are in blue and lysosomes are in orange. D shows segmented counts by frame, and E shows mitochondrial shape-category counts by frame, and F shows repeated detections. There are 2,084 frame-object observations of mitochondria and 2,344 of lysosomes. Display scaling does not change measurement inputs.

### Controlled and public image evaluation

For the five baseline synthetic sequences, mean pixel Dice was 0.967 for the revised default and 0.909 for the historical baseline. For the 50 sequences, these means dropped to 0.532 and 0.407, respectively (Figure 5). For the revised default, the mean link precision and recall were 0.422 and 0.468, respectively. Excluding ambiguous links increased accepted-link precision to 0.693 (defined in 34 of 50 sequences) and decreased recall to 0.178 across all 50 sequences. Mean identity switch counts were 0.18 and 15.34 for the revised default and historical exported-ID baseline, respectively. However, detection coverage was different in the historical baseline. Parameters and oracle-mask results indicate contribution of segmentation and distance gating in overall results. Results support a restricted operating range and do not show reliable tracking in crowded and/or low signal images.

Results for the seven held-out MoDL patches showed mean Dice was 0.228 for the historical baseline and 0.231 for the revised default. For the fixed top-hat configuration and fixed local thresholding, mean Dice was 0.421 and 0.395, respectively (Figure 6). The revised default and historical baselines both failed to recover foreground overlap in one of the MoDL patches. Results suggest that the default threshold alone is not adequate for this data. Of the 520 evaluations of the method for the four confocal studies, all 520 completed without execution errors. Legacy masks in those data contributed 12,505 object measurements. The original circularity formula exceeded one in 7,167 cases, and the definition of matched contours exceeded one in zero cases. Numerical geometry, not boundary accuracy, was assessed in this same-mask comparison. The confocal recordings supplied no independent reference trajectories, so execution of these commands does not prove tracking fidelity.

All 120 additional method-image evaluations were completed after the optional imagecodecs TIFF decoder was installed. The first attempt stopped at an LZW-compressed TIFF due to the absence of this dependency, and its log is retained. Mean Dice overlaps for the 13 Myofibril patches for the historical baseline were 0.039, 0.042 for the revised default, 0.039 with top-hat, and 0.047 for local thresholding (Figure 7A). The low Dice overlaps make the use of the settings, as is, not amenable for the annotated object interiors. It was noted that thresholding captured membrane fragments, and the reference masks included mitochondrial interiors. These representations reflect a plausible interpretation for the observed discrepancy and not a validation for a corrective algorithm. Local thresholding resulted in mean 22,271 components for the 17 Mito Hacker fields, as compared to 521 for the revised default (Figure 7B). Without reference masks, these counts indicate parameter sensitivity and do not validate the correct object number. No settings were retuned after scoring. The 13 annotated patches do not represent independent biological replicates.

## Biological tracking and fragmentation

The revised default matched 252 of 5,258 manual localizations and recovered 103 of 4,959 reference links after ambiguity exclusion. It achieved a mean per-movie localization recall of 0.067 and a mean accepted-link recall of 0.026. With fixed top-hat preprocessing, these means improved to 0.273 and 0.183, respectively, and 852 accepted reference links were recovered. The default entirely missed 227 of 284 trajectories and fully recovered none. The top-hat setting entirely missed 129 trajectories and fully recovered three. The default achieved an accepted-link recall ranging from 0.021 to 0.039 for the 2 to 5 pixel matching tolerances. These results show a poor transfer of the default segmentation to the vesicle movies. Conditional precision was 1.000 for the default among 103 accepted links with two reference-matched endpoints. Most predicted links were unverified. That restricted value does not prove overall tracking accuracy. Supplementary Figure 9 and Tables 6 and 7 show the results at the movie-level and counts of fragmentation.

## Particle refinement and reserved-movie evaluation

The baseline analyses are in Figures 2, 3, and 5 to 17 and Supplementary Figures 1 to 9. Particle detector and association rule refinements are in Supplementary Figure 10 and Table 9. Particle detection and association development used six real and nine synthetic movies and three noise-only controls. Evaluation used six reserved biological movies, 30 new synthetic movies, and five noise-only controls. We reran the original 50 synthetic sequences and 20 external annotated patches without further parameter adjustment. The static image evaluation used the selected per-frame detector without any temporal filtering.

**Figure 9.**
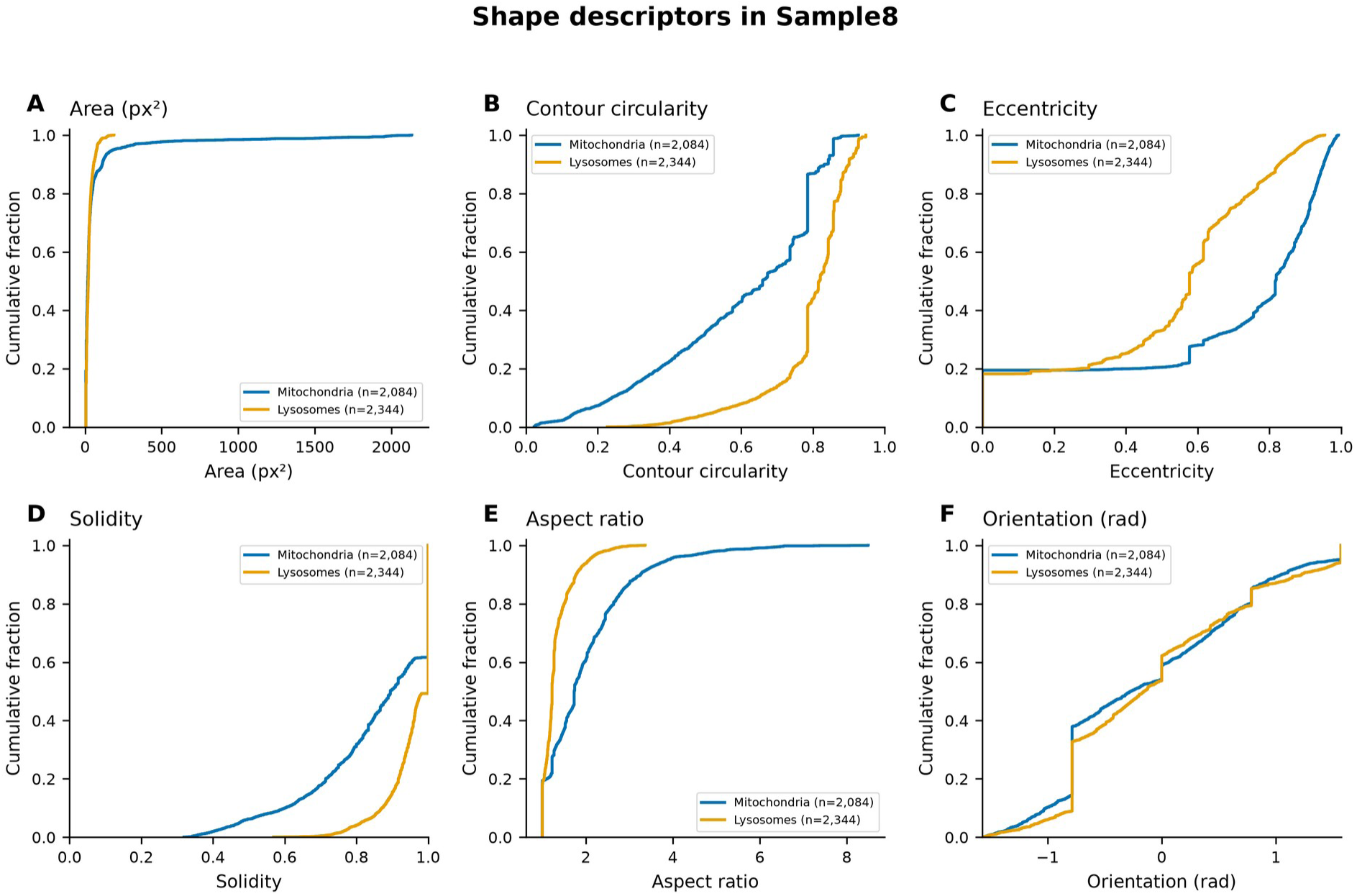
Shape descriptors. **A** through F show empirical cumulative distributions across 40 frames of pixel-count area, contour circularity, eccentricity, solidity, aspect ratio and orientation, respectively, for Sample8. Each legend shows the number of finite observations for a given descriptor. Circularity uses matched contour area and perimeter, and does not contain clipping. Plots show raw measurements without kernel smoothing.

Six reserved movies contain 145 annotated trajectories and 2,764 adjacent reference links. Mean accepted-link recall for the frozen default and fixed top-hat configurations was 0.014 and 0.212, respectively. With the particle pipeline, mean accepted-link recall was 0.696, and mean localization recall was 0.768. The new pipeline recovered 2,024 accepted reference links, and four additional accepted links related to different annotated identities, giving conditional precision of 2,024/2,028 among the links with two reference-matched endpoints. Remaining predictions were left unverified. Thirteen trajectories were completely missed, and 33 were fully recovered with accepted links, compared to 41 missed and two fully recovered with the top-hat configuration. Accepted-link recall was 0.739 with a 5-pixel tolerance and 0.587 with a 2-pixel tolerance. Individual reserved movies yielded accepted-link recall from 0.408 to 0.911. Supplementary Fig. 10 and Table 9 show the paired comparisons and their shortcomings.

The new detector together with the previous 6-pixel tracker yielded a mean accepted-link recall of 0.619 on the reserved movies. Replacing the association rule improved recall to 0.696 under the same three-observation requirement. Detection thus mainly drove the improvement, with association adding a smaller gain. Removing the length requirement improved recall to 0.707, but retained a higher number of noise detections. In 30 new synthetic movies spanning two motion patterns, three signal to noise ratios and five random seeds, mean detection precision was 0.967, accepted-link precision was 0.970 and accepted-link recall was 0.803. Five separate noise only movies retained an average of 0.34 detections per frame, as compared to 17.19 prior to the length filter. Analyst-defined controls provide complete references to assess false detections; they do not address biological specificity.

The original 50 synthetic sequences, when rerun, yielded mean Dice scores of 0.877, 0.752 and 0.532, for the new particle masks, the fixed top-hat configuration and the frozen default, respectively. Mean instance precision and accepted-link recall increased from 0.407 to 0.966 and 0.178 to 0.965, respectively. The two baselines reproduced their earlier results to within an absolute error of 10^-12. Transfer to the external mitochondrial masks was limited. Without the length filter, the selected detector yielded mean Dice scores of 0.369 and 0.060 for the seven MoDL SIM patches and the 13 Myofibril patches, respectively. The MoDL score was still below the earlier top-hat result of 0.421. The particle pipeline thus does not replace target-specific segmentation validation for elongated or densely packed mitochondria.

Mean fragmentation counts per sequence for the historical exported-ID baseline and the revised default were 21.80 and 3.76, respectively. With the exclusion of ambiguous links, mean fragmentation count for the revised default rose to 33.72, which is consistent with its limited accepted-link recall. The fixed 6-pixel gate provided 1.96 fragments and 3.54 accepted-link fragments per sequence. The default entirely missed a mean of 1.16 reference trajectories per sequence. Reporting fragmentation alongside coverage prevents low detection from appearing to be good track continuity.

### Core integration verification

The example adapter was written with LLM assistance and executed as traditional code without connecting an LLM to the analysis runtime. Guarded and direct calls agreed exactly across the 19 CSV and TIFF outputs of a known-motion fixture of 4 frames and the 20 corresponding outputs of the 40-frame Sample8 execution. The fixture maintained the expected speed of 1 px/frame. Comparison with the retained Sample8 results showed a maximum absolute numeric difference of 8.88 × 10⁻¹⁶. This is within the absolute tolerance of 10⁻¹². Eight deliberate mismatch controls were rejected before analysis, including an altered installed-package copy. An environment opt-in allowed different versions of NumPy and pandas, but still rejected the altered core.

The open Gemini desktop application generated an external adapter from the provided interface, and a fixed synthetic test contract. The application could not read or execute local files; therefore, the exact captured source was executed unmodified in the installed-wheel environment. Its 19 numerical CSV and mask or label TIFF outputs matched standalone command line outputs, and reported a maximum numerical difference of 0. Both channels returned one object per frame over four frames and a mean speed of 1 pixel per frame. The notebook route also successfully executed all three code cells. These checks establish integration on the supplied fixture, without establishing accuracy on new images.

The original baseline release included 51 local software tests. Results from known-motion analyses were retained. The 2.5.0 release adds 21 tests for particle detection and association, temporal filtering, and output integrity and consistency. The Sample8 input was processed through the particle route in pixel and frame units. That process does not replace the baseline figures, and has no manual organelle reference. Therefore, it does not prove improved accuracy of neuronal analyses.

For baseline analysis of Sample8, the median wall time without PNG rendering was 2.21 seconds, with a range of 2.21 to 3.02 seconds. The median peak resident memory was 336.4 MiB. With PNG rendering, the median time was 2.42 seconds (with a range of 2.42 to 2.67 seconds) and the median peak resident memory was 410.3 MiB. These timing results are for a 40-frame, two-channel, 258 x 258 input on the recorded host; they are not estimates for scaling with larger movies.

AutoMorphoTrack 2.5.0 passed 72 tests in each of 12 GitHub CI installations. It built wheels and source distributions for Windows, Linux, and macOS, with Python 3.11 and 3.12. All installations kept the 18 baseline and five particle source modules. Each installation compared 19 baseline and 19 particle CSV or TIFFs with recorded references. Integer masks were exact, and for baseline and particle outputs, numerical tolerances were 10^-12 and 10^-9, respectively. The test failed initially on Windows due to Git line-ending conversion which changed the source hashes. Enforcing LF line endings solved this, and the numerical code remained the same. The OpenCV import failure and its system-library remedy documented for minimal Linux remain documented.

### Public image workflow and quality control

Segmentation and correspondence for early endosomes are shown in Figure 2. The first six full fields contained 49 accepted links, and 750 flagged, ambiguous assignments under the fixed 15-pixel gate. Figure 3 shows the result of the separate alignment check for a known translation of one public frame. All five tested integer shifts were exactly recovered. Normalized RMSE over the 58,560 valid pixels, for the displayed shift, changed from 0.1057 before alignment to zero afterward. This controlled translation test is separate from the primary tracking measurements.

### Detection, Counting, and Morphological Classification

The segmentation of Sample8 produced 2,084 mitochondrial and 2,344 lysosomal frame-object detections across 40 frames **(Figure 8)**. These totals include repeated observations. The frames contained 44 to 61 mitochondrial components and 51 to 65 lysosomal components. The blue and orange overlays show the respective masks, and the scale bar indicates a distance of 50 px.

The mitochondrial categories in Figure 8E and F summarize eccentricity-based shape descriptions, including an intermediate category. They do not describe fragmentation, fission, or fusion.

### Shape Profiling and Feature Extraction

Figure 9 shows the empirical cumulative distributions of area, contour circularity, eccentricity, solidity, aspect ratio, and orientation. Each panel indicates the number of finite frame-object observations. The revised contour geometry and bounded plotting replace the original inconsistent circularity values and smoothed distribution tails.

### Organelle Tracking and Motility Analysis

In Sample8, 79 mitochondrial and 73 lysosomal candidate tracks contained accepted motion steps **(Figure 10)**. The corresponding motion summaries used 333 and 524 accepted adjacent-frame links, respectively. Ambiguous mitochondrial and lysosomal links totaled 1,324 and 1,574, respectively, and were ignored. Accepted links represented 20.1% of assigned mitochondrial links and 25.0% of assigned lysosomal links. These percentages describe retained coverage. Therefore, the distributions describe a retained subset. Path length and mean speed satisfy the relation path length = mean speed × accepted duration, which explains structured bounds in the scatter plot of the mean speed and path length of mitochondrial and lysosomal candidate tracks, without implying a biological threshold of minimal movement.

**Figure 10.**
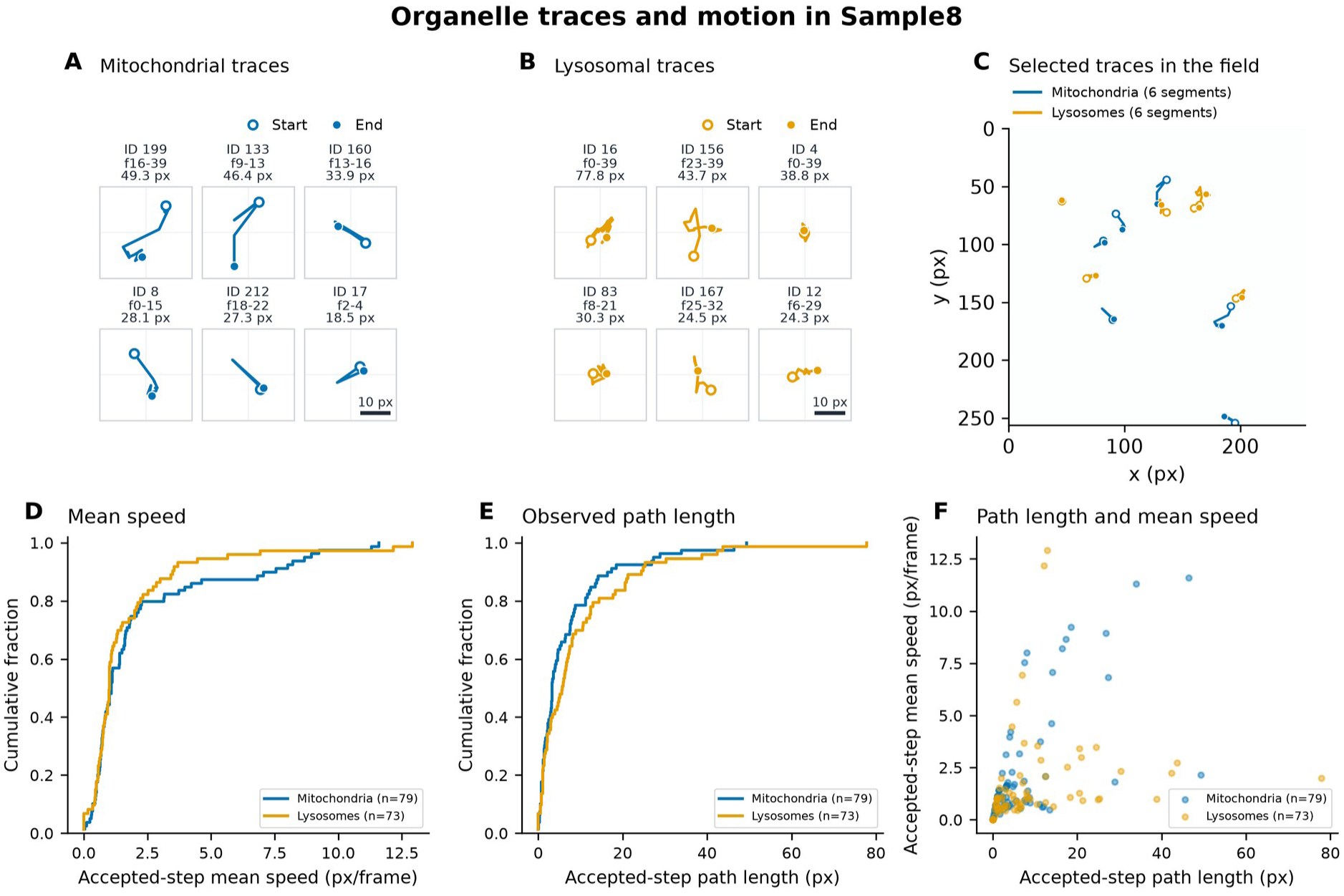
Organelle traces and motion summaries. **A** and B display the six continuous accepted trajectory segments with the longest path length for mitochondria and lysosomes, respectively. Each contains at least two accepted steps; gaps or ambiguous links end a segment. Traces are centered in 30-pixel-wide views at the same scale with no smoothing or rotation. Markers identify the beginning and end; labels provide track ID, zero-based frame range and path length. C illustrates the selected segments at their original image coordinates. Examples have been selected to illustrate movement; D-F retain all finite movement summaries. D shows accepted step mean speed, E shows observed path length, and F plots the related quantities. Summaries are finite, and include 79 mitochondrial and 73 lysosomal tracks, and 333 and 524 accepted links respectively. 1,324 and 1,574 ambiguous links were ignored, respectively. Observed path lengths include only retained steps, and do not have to be continuous, and speeds are in px/frame. Distances are in px.

### Mask overlap and intensity descriptors

Figure 11 distinguishes mask overlap from native intensity measurements. The mean Jaccard overlap and mean intensity Pearson correlation for the 40 Sample8 frames were 0.0285 and 0.1837, respectively. The mask conditioned mitochondrial and lysosomal intensity fractions averaged 0.0341 and 0.0722, respectively. These are descriptive measurements within the full crop and do not establish physical contact of organelles.

**Figure 11.**
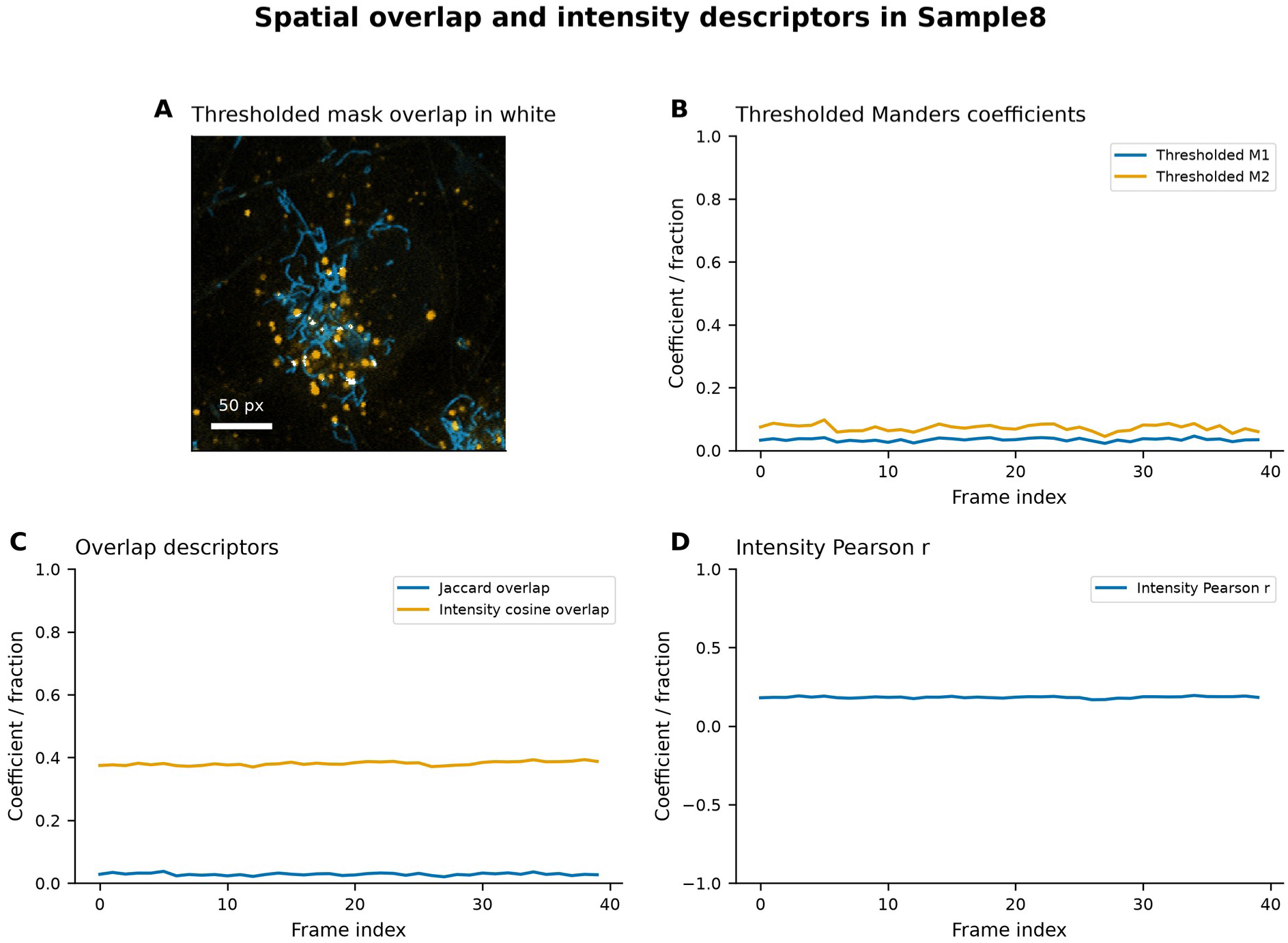
Mask overlap and intensity descriptors. A shows native-intensity channels with mask intersection in white at frame 0, and a scale bar at 50 px. B shows mask-conditioned intensity fractions. C shows Jaccard mask overlap and intensity cosine overlap, and D shows native-intensity Pearson correlation. Each time series is composed of 40 frame measurements from the same full-crop analysis region. Missing coefficients are undefined. Image overlap does not prove membrane contact or functional interaction.

### Integrated Correlation Summary

Figure 12 limits each frame to one summary and removes identifiers. The Spearman matrix describes associated frame summaries, and pairwise valid-frame counts are provided in the source data. Biological meaning is limited due to temporal dependence and shared metric definitions.

**Figure 12.**
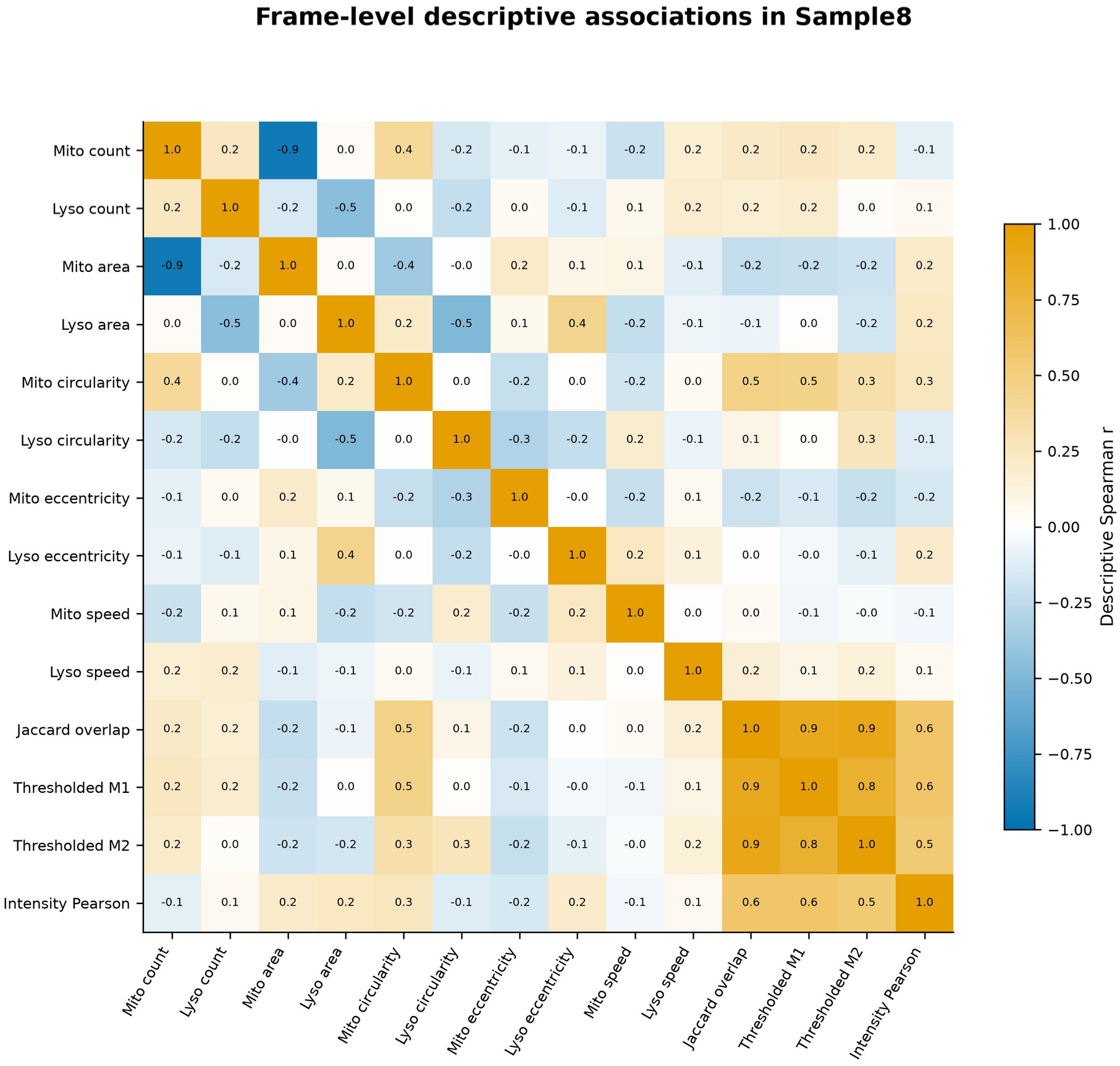
Descriptive frame correlations. Spearman correlations of frame-wise means of shapes, accepted-step speeds, counts and colocalization metrics in Sample8. There is one row per frame, for up to 40 paired observations. Identifiers are omitted and missing data are handled pairwise. Exact paired frame counts are supplied in Frame_Correlation_N.csv. The matrix is descriptive and is temporally dependent; no hypothesis tests are performed.

### Descriptive region comparisons

Sample6 and Sample8 illustrate comparisons of counts, shape descriptors, accepted motion summaries, and image colocalization. Outputs complementing these comparisons are shown in Figures 13 to 17, with the configuration and source measurements retained for each region.

**Figure 13.**
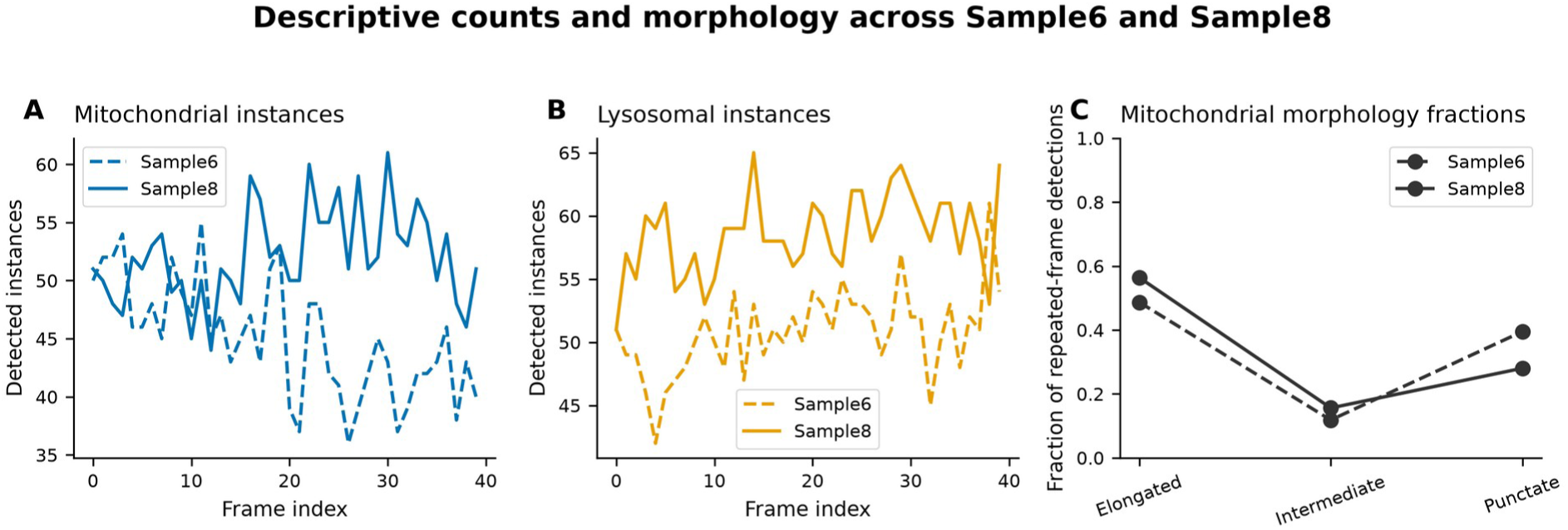
Descriptive region counts and morphology. **A** and B present frame-level counts of mitochondria and lysosomes in Sample6 and Sample8. C presents shape-category fractions of mitochondria pooled from repeated frame-level detections. Each region consists of 40 frames. Sample8 is represented by solid lines; Sample6, by dashed lines.

Figure 13 shows shape-category fractions and counts for Sample6 and Sample8. Each region consists of 40 frames captured from the same video. The differences shown are descriptive and do not represent analyses of independent cell or treatment effects.

Figure 14 shows the same six shape descriptors as in Figure 9. The color of the line represents the organelle of interest, and the line style represents the region. Repeated detections across frames contribute to the plotted distributions.

**Figure 14.**
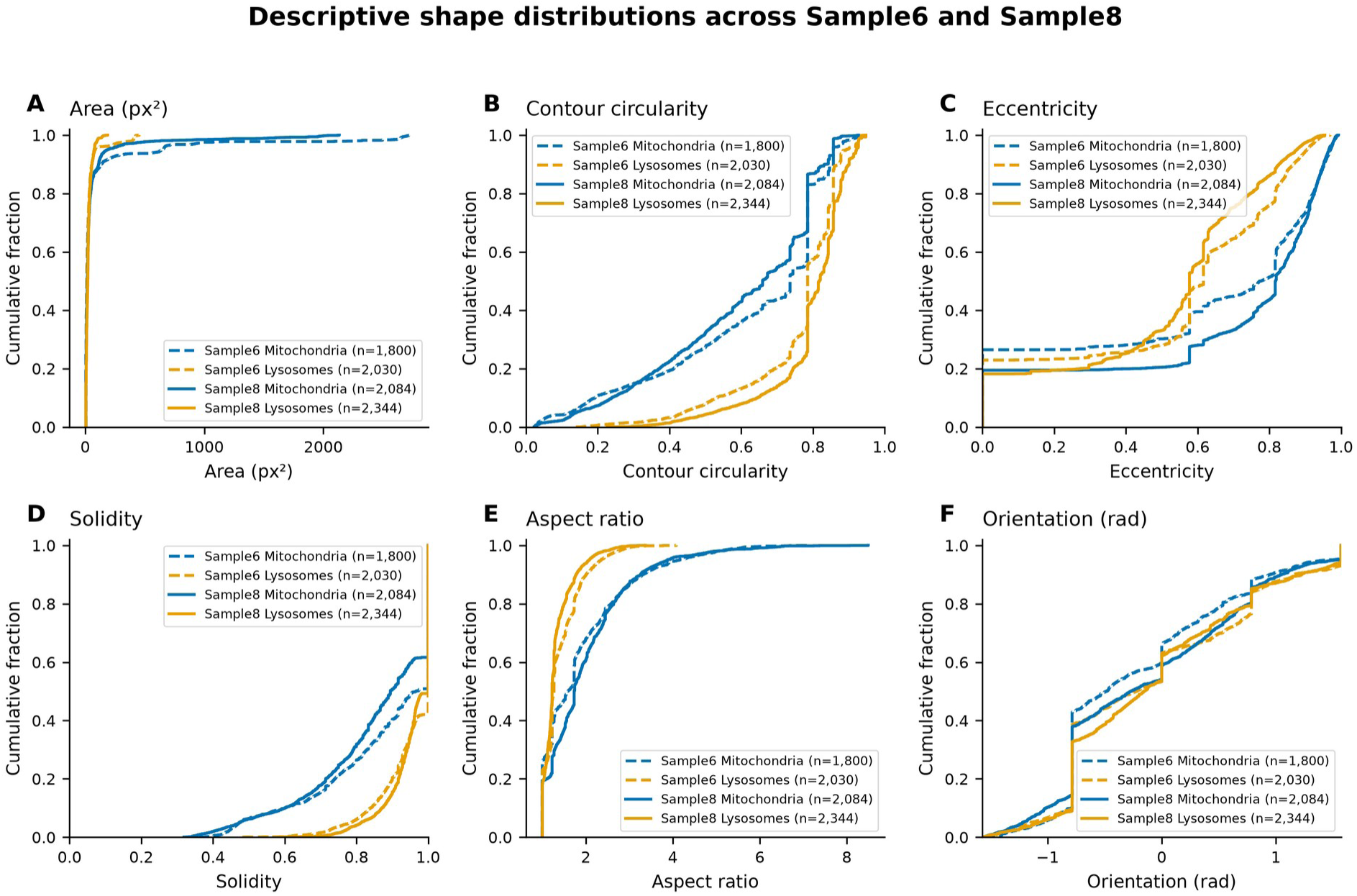
Descriptive region shape distributions. A through F analyze the pixel-count area, contour circularity, eccentricity, solidity, aspect ratio, and orientation of Sample6 and Sample8. Mitochondria and lysosomes are represented by blue and orange, respectively. Line styles identify regions. Each empirical cumulative curve states the number of finite frame object observations. Each region consists of 40 frames.

Fig. 15 shows accepted step mean speed and observed path length for each channel. Sample6 contributed 94 mitochondrial and 74 lysosomal tracks with finite motion summaries; Sample8 contributed 79 and 73. Ambiguous links place limits on which motion contributes to these distributions.

**Figure 15.**
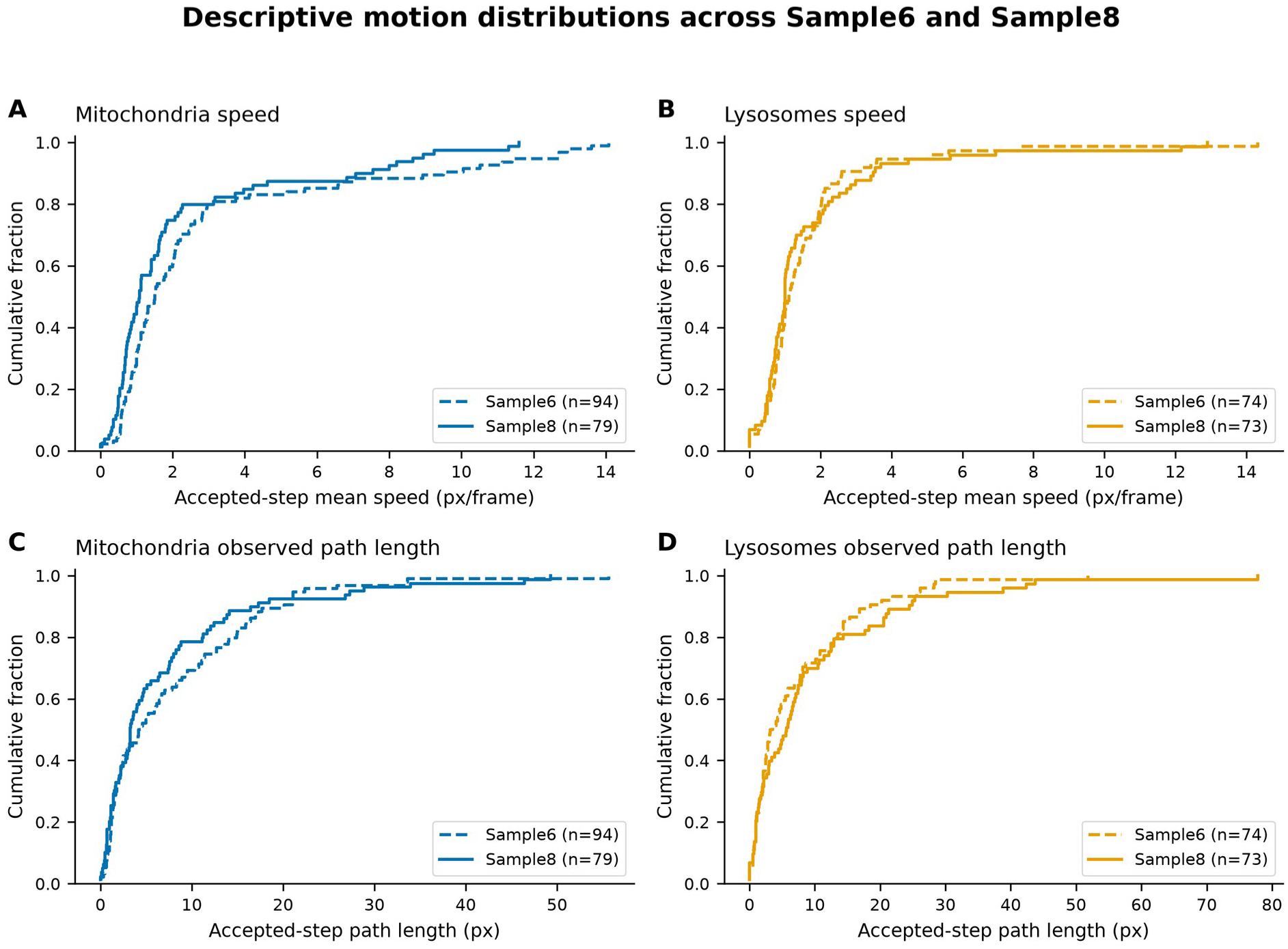
Descriptive region motion distributions. A and B analyze mean speeds of the accepted steps for mitochondria and lysosomes, respectively, and C and D analyze the corresponding accepted path lengths. There are 94 mitochondrial and 74 lysosomal tracks with finite summaries in Sample6, and 79 mitochondrial and 73 lysosomal tracks with finite summaries in Sample8. Distances are in pixels and speeds are in pixels per frame. Only accepted adjacent steps are analyzed; ambiguous steps are ignored.

Fig. 16 shows the two mask-conditioned intensity fractions, intensity Pearson correlation, and Jaccard mask overlap, for the 40 frames in each region. These assess optical and intensity overlap and association at the resolution of the data collected.

**Figure 16.**
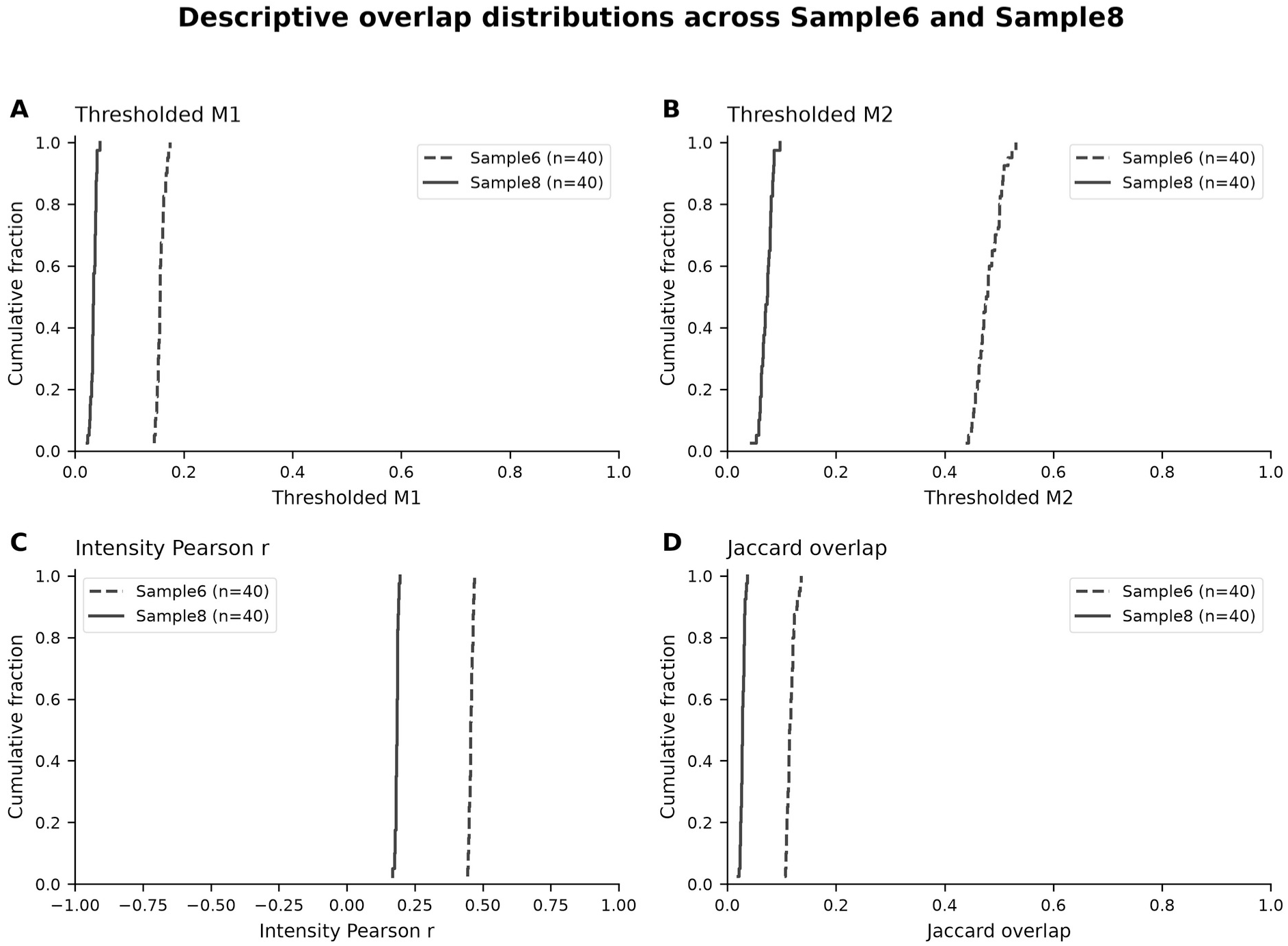
Descriptive region overlap distributions. A through D evaluate mask-conditioned intensity fractions of mitochondria and lysosomes, Pearson correlation of native intensities, and Jaccard mask overlap, respectively. The full range of Pearson correlation from −1 to 1 is retained. Descriptors of Jaccard mask overlap are in the range [0, 1]. Each region contains 40 frame measurements.

Fig. 17 shows the Sample6 Spearman correlation matrix subtracted from the Sample8 matrix. The color scale is from −2 to 2. The source matrices describe temporally dependent frame summaries; their difference is not a significance test, and does not indicate changed biological coordination.

**Figure 17.**
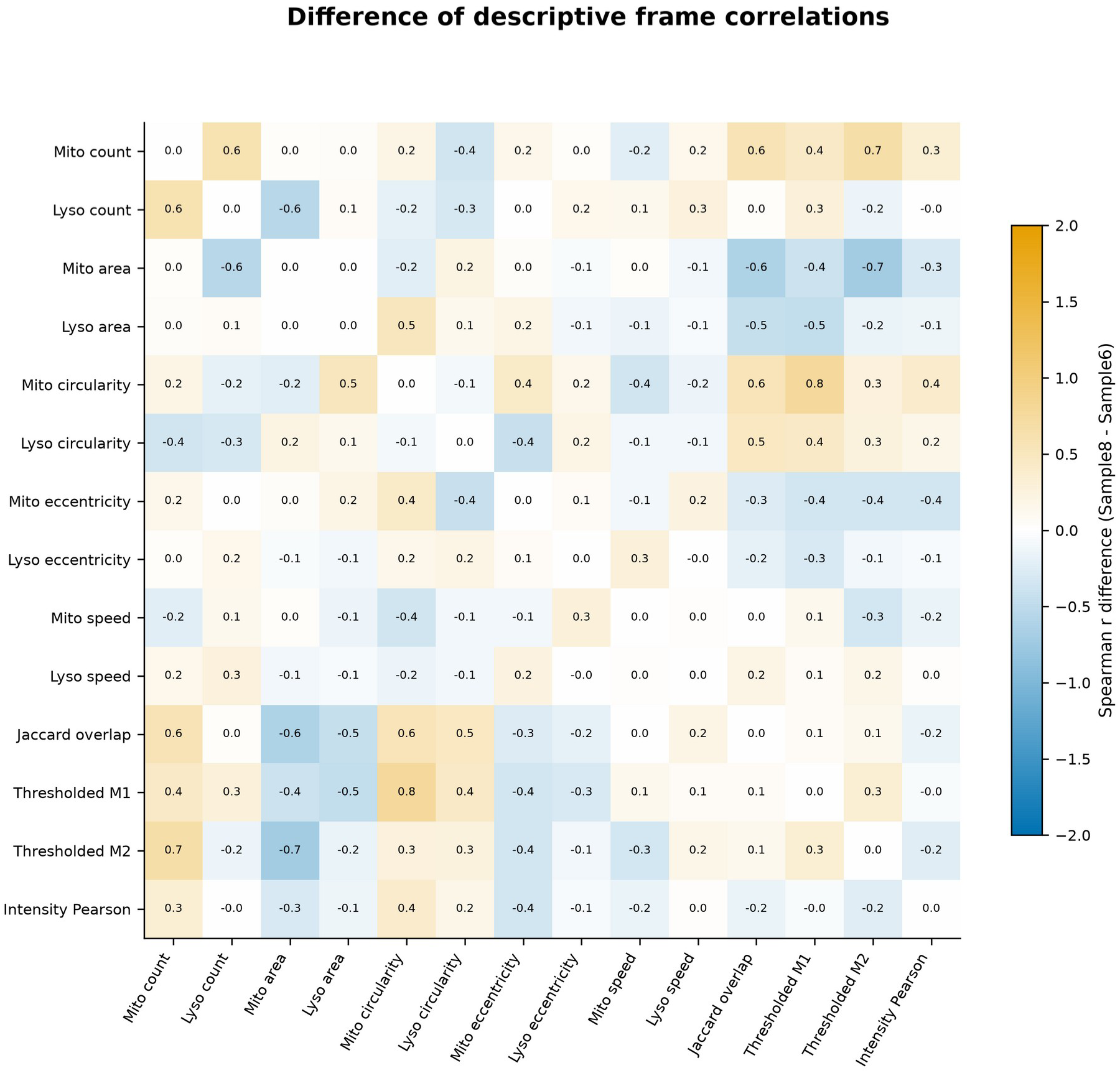
Difference of descriptive correlations. Each entry is the Sample8 Spearman coefficient minus the Sample6 coefficient, for the same pair of frame summaries. The color scale is from −2 to 2. Both source matrices use up to 40 temporally dependent frames, and the exact counts of valid pairs are provided separately. This subtraction is for description, not for statistical inference.

## Discussion

AutoMorphoTrack links established segmentation, measurement, and tracking operations through shared masks, persistent identities, explicit units, and recorded configuration. Its main contribution is the inspectable implementation of this workflow. Comparisons with other tools describe its scope, and accuracy experiments compare the historical AutoMorphoTrack implementation with fixed revised configurations. External segmentation models and language-model applications retain their own scope, capabilities, and limitations.

Prior work resulted in image output, but the quantitative accuracy of those images was not established. This work fixes defects in object linkage and geometry, and metric definitions. It evaluates the behavior against controlled standards. Performance depends on the quality of the inputs and on the operating range. A successful run, or a visually plausible output, is not proof that the output segmentation and tracks are correct.

Segmentation performance varied with source images and annotation target. Top-hat correction and local thresholding improved the agreement with the held-out MoDL masks. However, all four of the settings tested gave low overlap with the Myofibril reference masks. The membrane label and the filled object annotations help explain the failure of the bright-pixel segmentation in this setting. The improved package accepts external instance labels, which enables the use of other segmentation methods. The interface, however, does not guarantee the accuracy of the methods. These results support the need for a reference-based evaluation for each case in view of the mask inspection.

Measurements of motility are contingent upon detection and correspondence across frames. Tracks can be lost or ambiguous links created due to crowding, object entry or exit from the field of view, touching other objects, or changing appearance. Persistent identifiers and 1-to-1 assignments resolve some defects in the original implementation of the software but do not resolve all tracking ambiguities. Image colocalization is a measurement that is limited by resolution. Therefore, even if image colocalization shows two puncta are in close spatial proximity to each other, it cannot prove membrane contact. Image colocalization by itself also cannot determine if two puncta are functionally coupled or if mitophagy has occurred.

The updated particle pipeline improved recovery of annotated vesicle trajectories. Parameters were set before evaluating reserved movies, which are part of the same, previously examined study. Generalization to other acquisition systems, organelle labels, or biological states remains unresolved. The three-observation requirement reduced noise, and in the case of short-lived events, may have eliminated them. High precision among reference-matched endpoints does not necessarily explain or characterize the many unannotated predictions. The external mitochondrial mask scores were low. Users should review retained and raw detections, set a segmentation target, and verify coverage prior to interpreting counts or motion in a biological context. The baseline set for the original analysis remains, and can be reused to reproduce that work.

Standardized exports facilitate downstream analysis especially when experimental sampling structure is retained. Frame-level Spearman correlations indicate relationship and dependency among the observed data, which can reflect temporal dependence or shared mathematical components, but do not describe mechanism. Comparison of experimental groups requires independent sampling and analysis which accounts for observations that are nested within cells and experiments.

The publicly available independent datasets expand the original scope of testing. Annotated data enable users to evaluate the accuracy of a system; unannotated data only allow users to check the functionality of the system, inspect quality, and evaluate the sensitivity of the system to various parameters. Results from static images do not validate tracking. Results from a different fluorescent structure do not establish mitochondrial or lysosomal accuracy. Performance claims are made based upon the distinctions identified.

Keeping bytes of packages identifies the implementation used for a run, but new images still require adequate segmentation and tracking evaluation. Changes to wrapper logic, channel selection, preprocessing, settings, or downstream statistics can alter inference, even when the core is unchanged. Quantitative core changes require a new package version and relevant revalidation.

## Conclusion

AutoMorphoTrack provides a user-editable Python workflow for the segmentation, shape measurement and tracking of organelles within cells and evaluation of colocalization of cellular components. The revision makes transparent object identities, metric definitions, calibration choices and analysis provenance. Its utility depends on the accuracy achieved for the images to which it is applied, plus inspection of masks and tracks associated with those images.

The demonstrated operating range is limited by controlled tests and public data sets. Other organelles, microscopy conditions and experimental comparisons require supplementary reference data and independent sampling. The workflow allows measurements to be saved for such studies, limited by the underlying images and algorithms.

## Supporting information

Validated Core and Visual Guide

## Code and Data Availability

The AutoMorphoTrack GitHub release v2.5.0 contains an updated package, notebook, benchmark scripts, validation records, source data tables, figures, and manuscript files (22). The accompanying reproduction archive records dataset licenses, selections and file hashes. This release supersedes earlier revision deposits; historical releases remain available for reproducing earlier analyses.

The pre-revision bioRxiv manuscript, version 5, is available at https://www.biorxiv.org/content/10.1101/2025.07.19.665650v5. The revised analyses in this manuscript replace the previous ones.

AutoMorphoTrack 2.5.0 includes the original benchmark implementation in automorphotrack.validated and a particle pipeline in automorphotrack.refined (22). The 18 baseline modules remain the same as in 1.1.0rc2, while the five particle modules contain a separate frozen source digest. Package versions indicate distributions, while source manifests indicate numerical implementations. Earlier 2.x interfaces remain out of scope for validation. The release includes the original wheel and execution records for historical analyses.

The GitHub release includes an executed notebook for AutoMorphoTrack 2.5.0, as well as installation instructions and an example of guarded integration that verifies both numerical namespaces.

Older notebooks and comparison scripts are retained for provenance. The revised comparisons remain descriptive and retain their sampling units. Public benchmarking sources and their licenses are described in Supplementary Table 2; exact file selections and hashes are provided in the data manifests.

## Author Contributions

**A.B.** conceptualized and developed AutoMorphoTrack, designed the computational architecture, implemented the image-analysis pipeline, and performed all experiments, analyses, and figure generation. **J.G.S.** contributed to the writing and editing of the manuscript. **X.C.** supervised the project.

## Supplementary Figures

**Supplementary Figure 1.**
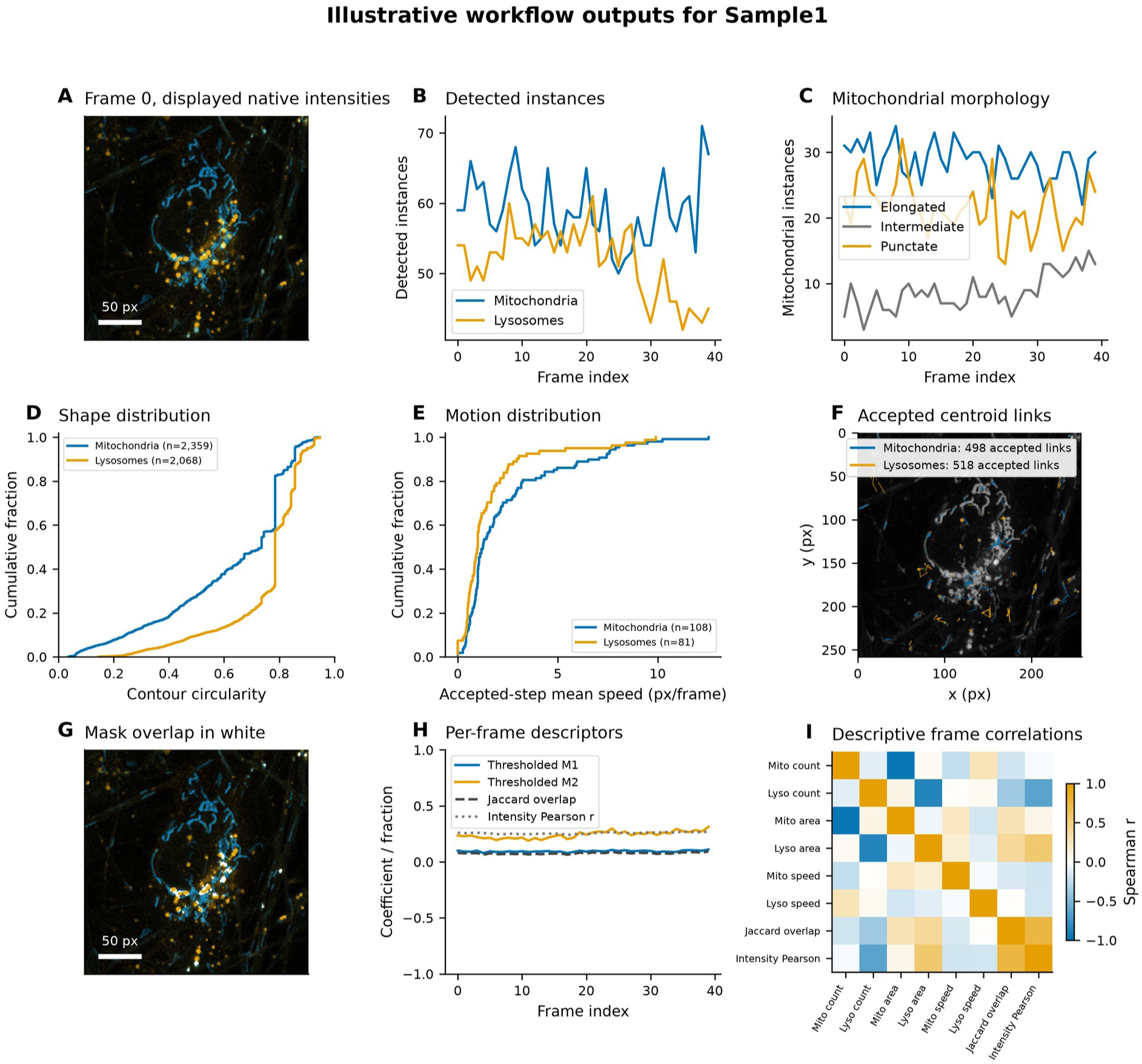

**Supplementary Figure 2.**
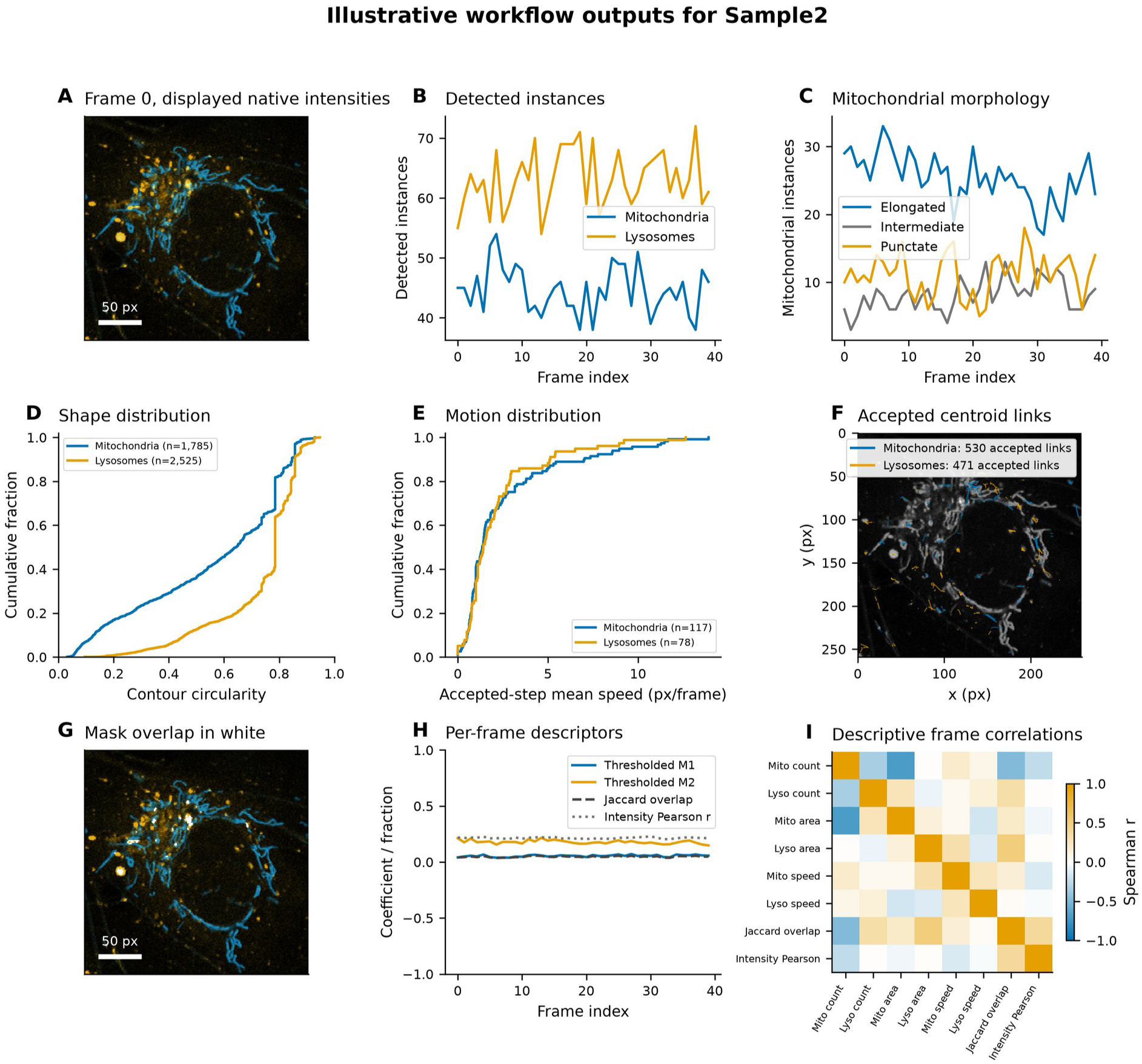

**Supplementary Figure 3.**
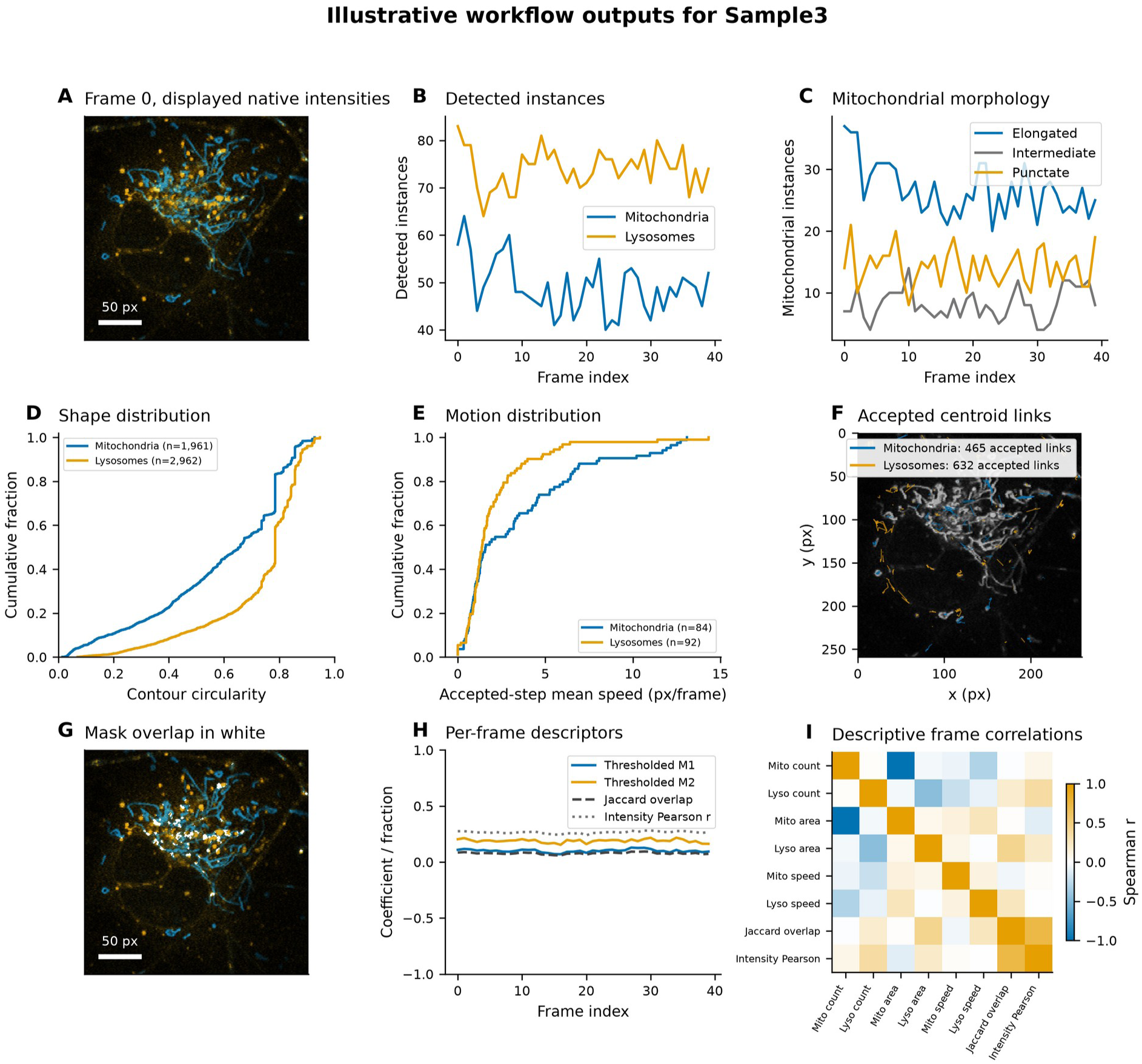

**Supplementary Figure 4.**
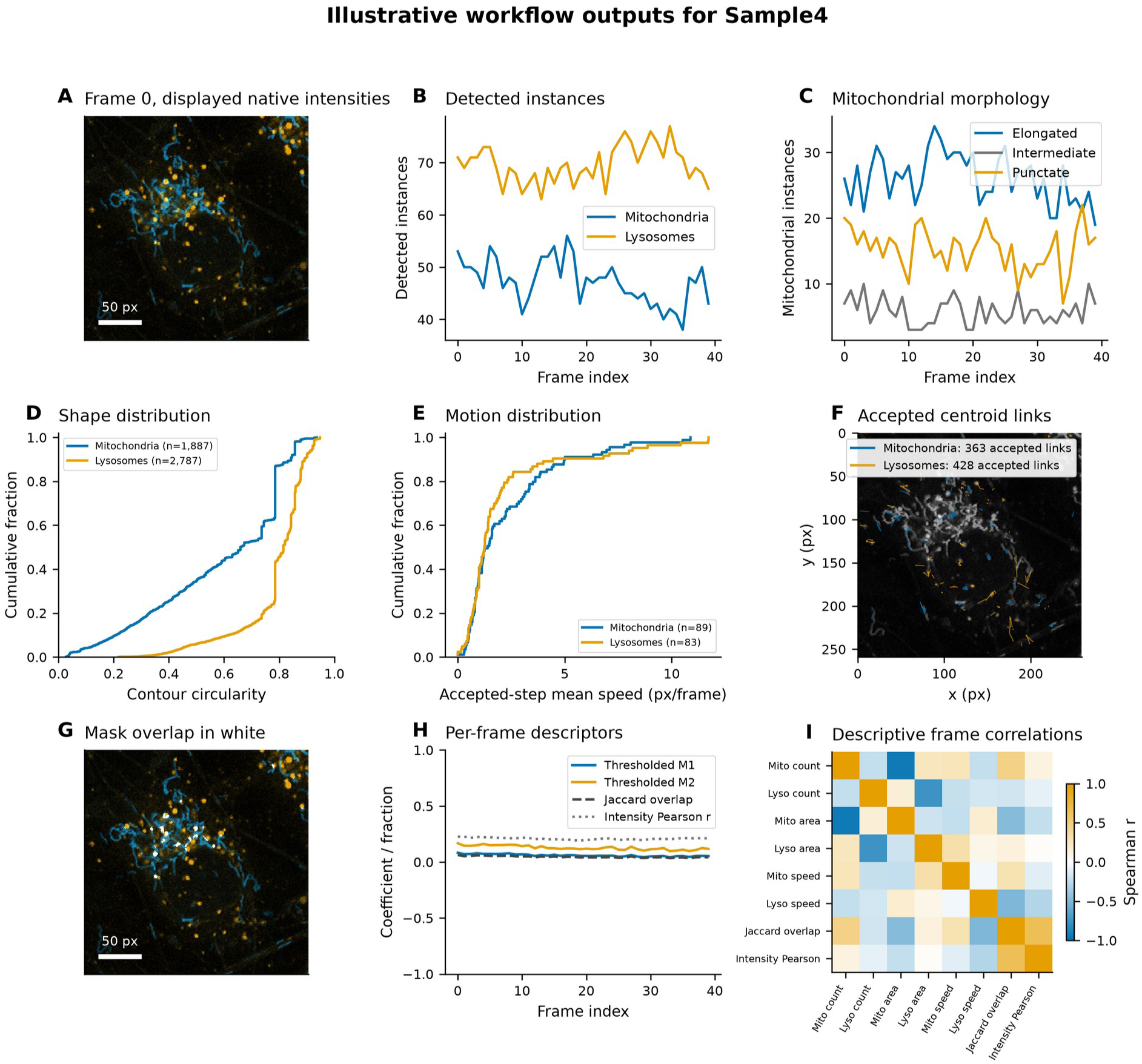

**Supplementary Figure 5.**
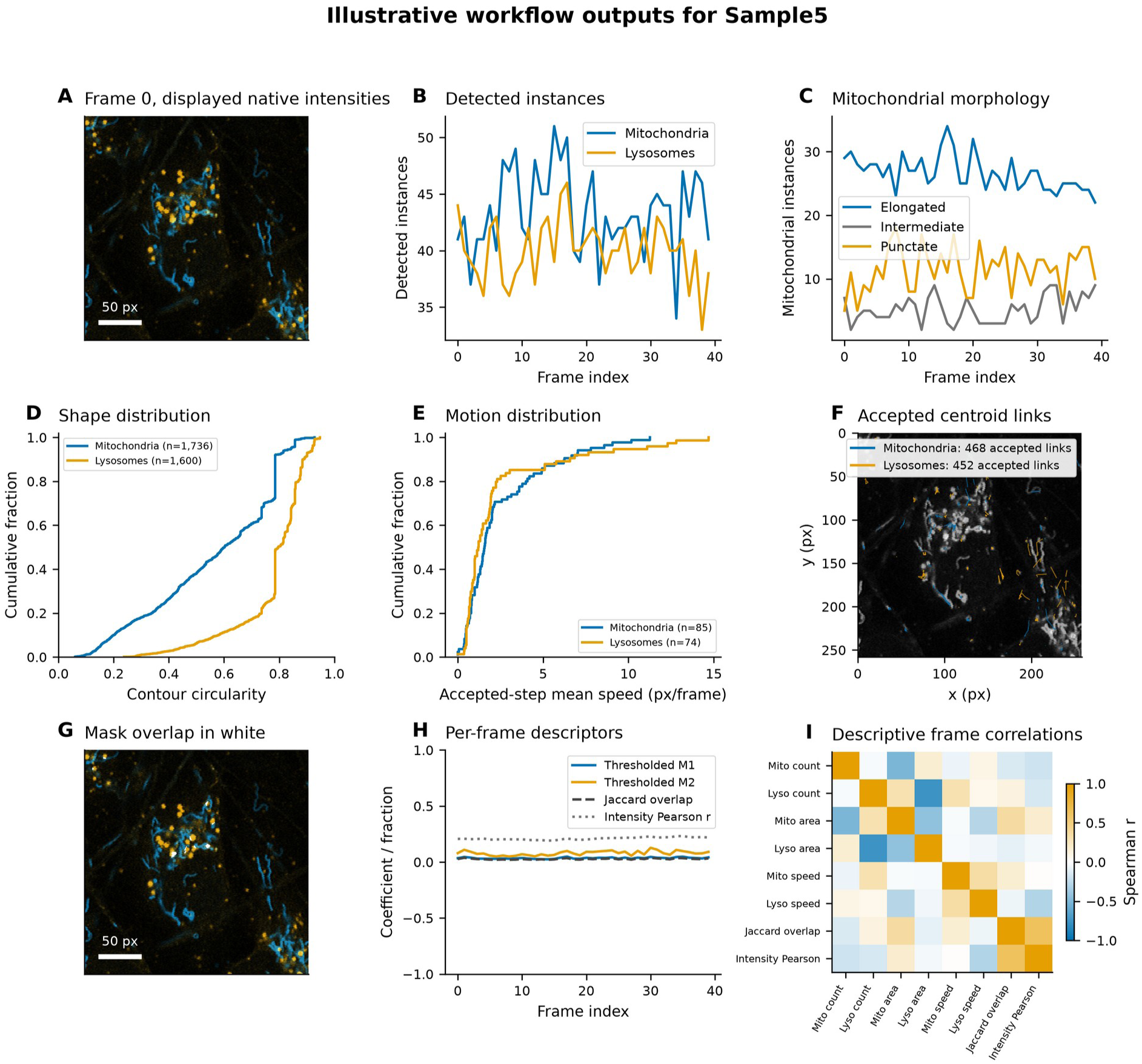

**Supplementary Figure 6.**
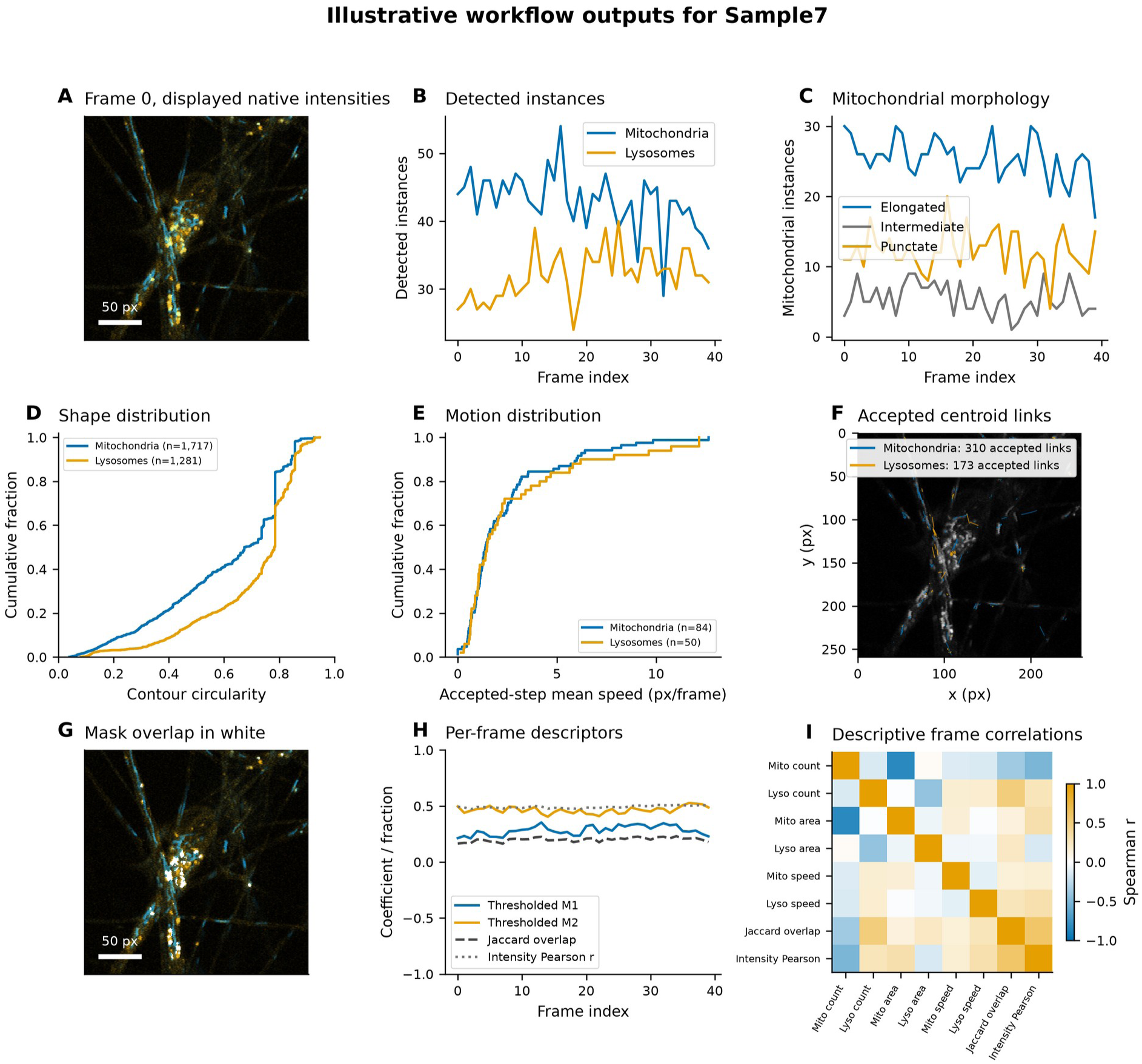

**Supplementary Figure 7.**
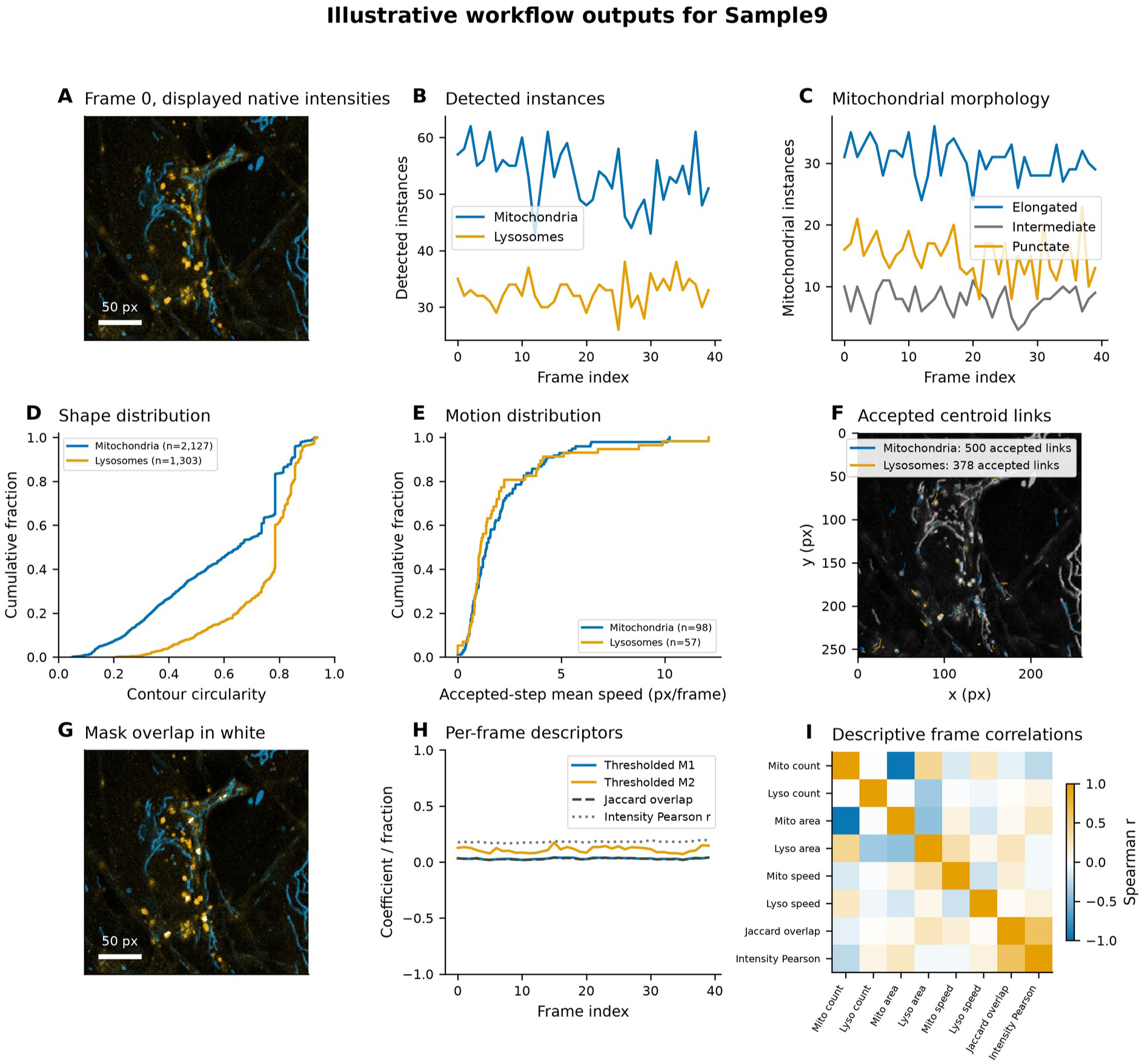

**Supplementary Figure 8.**
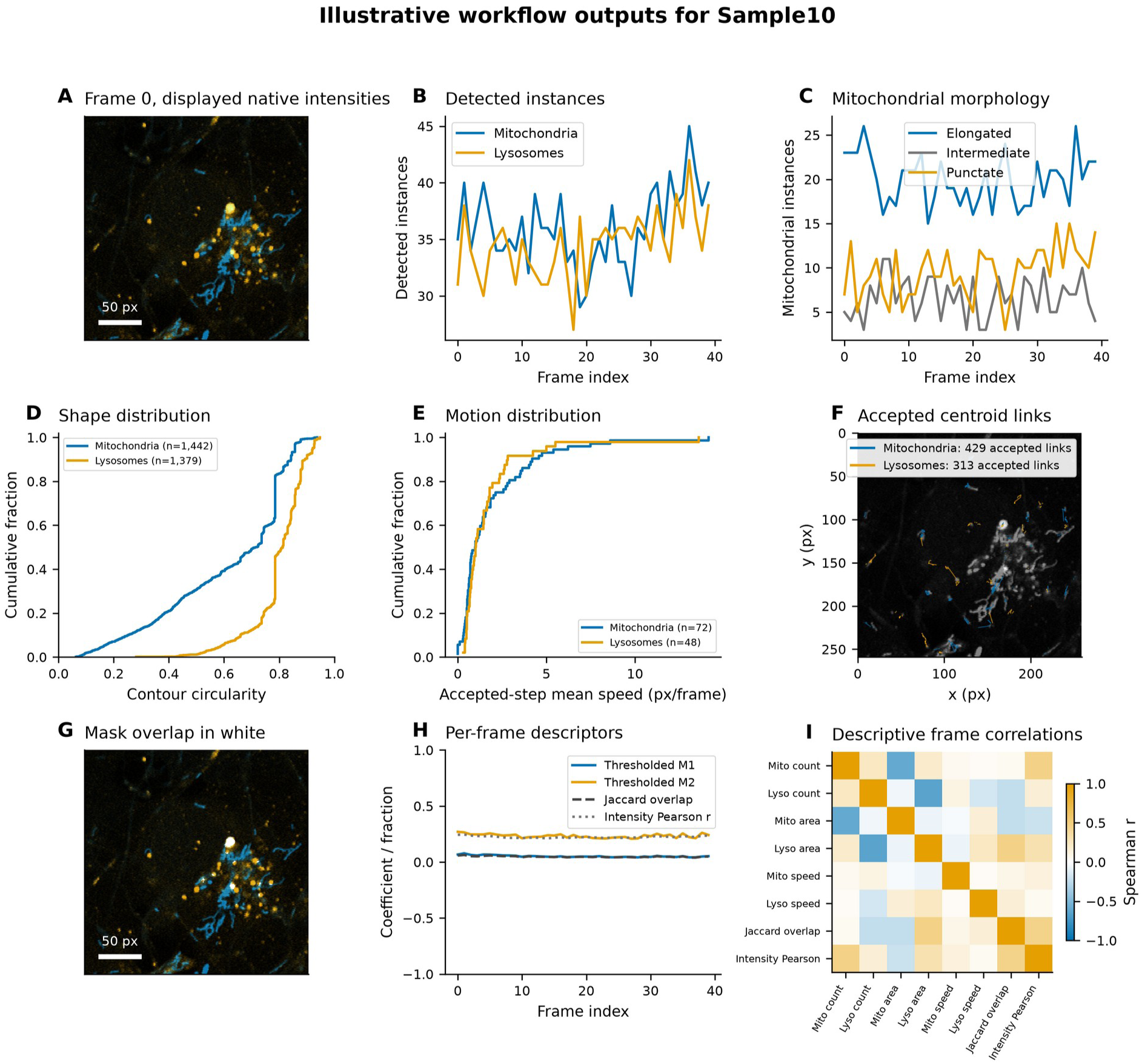

**Supplementary Figures 1 to 8 Additional image regions**

Figures S1-S8 show Sample1, Sample2, Sample3, Sample4, Sample5, Sample7, Sample9, and Sample10, respectively. Each region consists of 40 frames. A shows natively displayed frame intensities, B shows segmented counts, and C shows mitochondrial shape counts. D and E show contour circularity and mean speed with accepted steps, respectively, as empirical cumulative distributions with finite observation counts. F shows accepted centroid links. G and H show mask intersection in white, and overlap and intensity description, respectively. I shows descriptive frame-level Spearman correlations. Blue denotes mitochondria; orange denotes lysosomes. 50 pixels denote scale. Full run settings and per-region counts are shown in the source data.

**Supplementary Figure 9.**
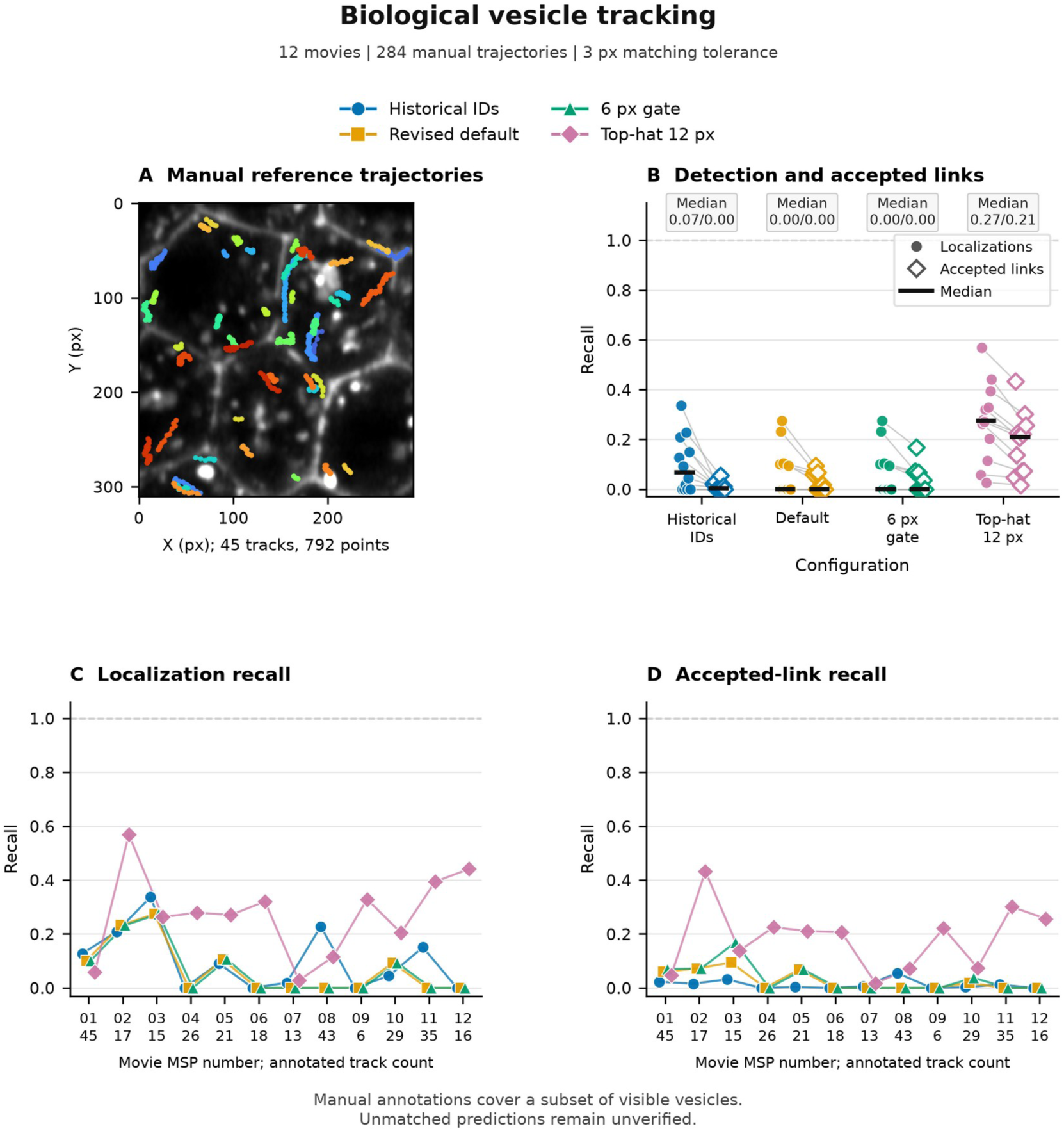
Biological vesicle tracking. Tracking of biological vesicles against manually deposited references. A shows frame 0 of the MSP01 Airyscan movie, along with the 45 manually deposited trajectories in their original coordinates. Lines stop at annotation gaps; paths connect annotated frames and do not indicate concurrent positions. Gray-scale images use the 1st and 99.5th percentiles of the intensity distribution. B shows the recall of localizations and accepted links for each of the fixed configurations for the 12 movies, and gives median values. C and D show individual movie values and annotated trajectory counts. Connecting lines identify configurations across movies and do not indicate a sequence in time. Matching predicted localizations to manual points uses a 3-pixel tolerance. The 284 manually annotated trajectories in secretory vesicles are used as a partial reference, and therefore predictions not matched with these references remain unverified. Image and manual annotations are from Richens et al. and the version 1 Cambridge deposit (23, 24), made available under a CC BY 4.0 license. The figure shows secretory vesicles.

**Supplementary Figure 10.**
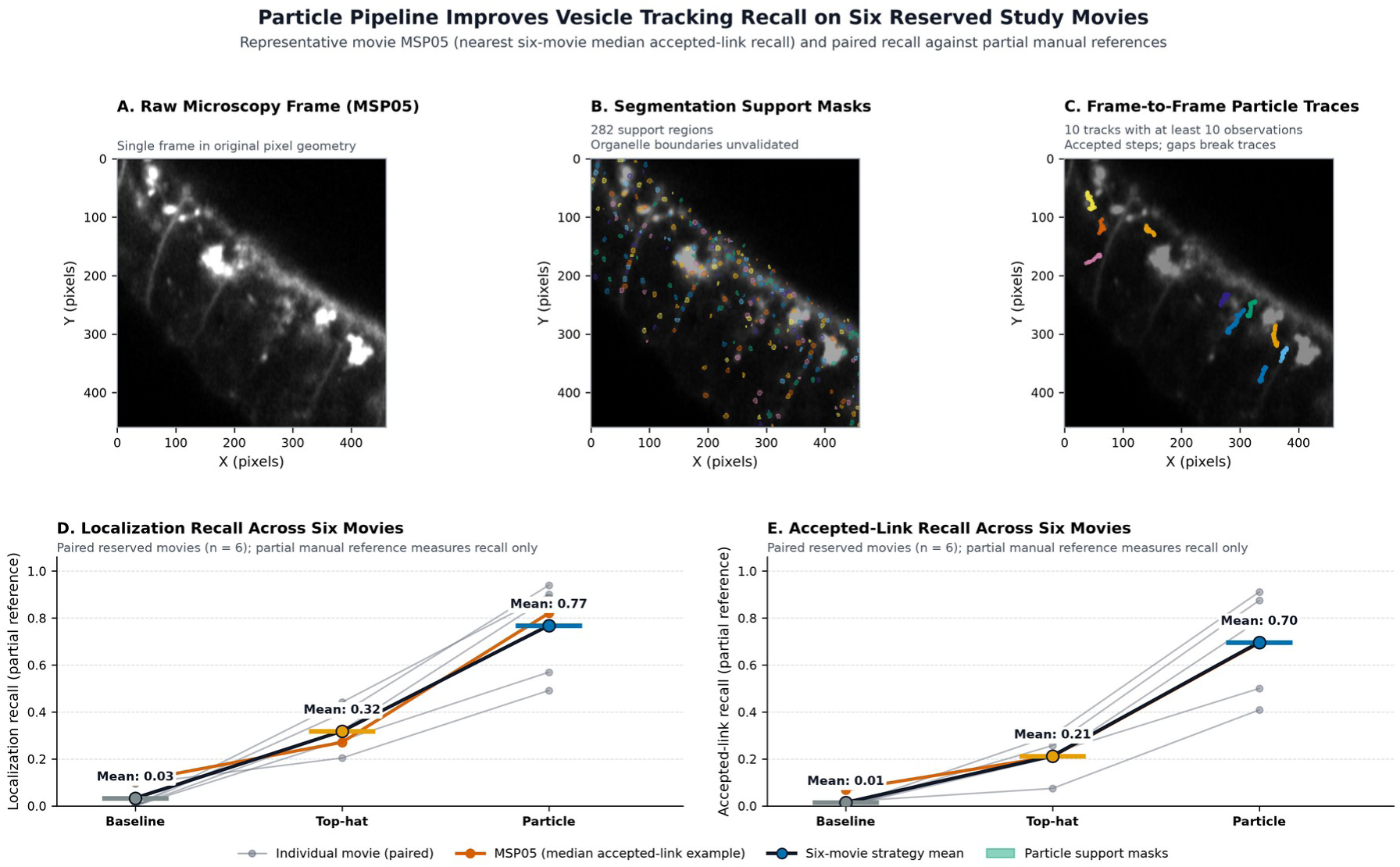
Particle segmentation and tracking. Segmentation and tracking of particles in reserved vesicle movies. Part A. Frame 0 of MSP05 from Richens et al. benchmark (23, 24). We selected MSP05 as the movie nearest to the median accepted-link recall of the six reserved movies; a tie with MSP06 was broken by sequence identifier. B. Overlays 282 retained support masks on this frame. Masks represent support fluorescence and have not been validated as organelle membrane boundaries. C. Shows 10 tracks with at least 10 observations and the largest first-to-last distance. Adjacent accepted steps are joined; gaps and ambiguous links break tracks. The image coordinate system is as is, and both axes are in pixels. D and E show a 3-pixel match tolerance for localization recall and accepted-link recall, respectively, for the six reserved movies. Thin lines connect measurements from the same movie, orange identifies MSP05, and large circles show the six-movie arithmetic means. The manual annotations cover selected vesicles, so these panels measure reference coverage and do not show overall precision. Display scaling does not change the absolute values of the data. Data DOI: 10.17863/CAM.114338, version 1; paper DOI: 10.1371/journal.pbio.3003099; CC BY 4.0.

### Supplementary Movie 1

Supplementary Movie 1. 40-frame Sample8 recording of native intensity. ROI with mitochondria in blue, lysosomes in orange. Image is an additive pseudocolor display and is not a mask-overlap display. Each channel uses its fixed first-to-99.5th percentile intensity limits calculated across all 40 frames. The annotation is a zero-based frame index. Playback is 5 frames per second for viewing only; it is not representative of the acquisition interval. The MP4 is an 8-bit lossy visualization, and is not a quantitative or raw-intensity data file. The associated Sample8_native16.ome.tif preserves the input uint16 pixels and channel order; the ROI is linked to the source capture of the file.

### Supplementary Movie 2

Supplementary Movie 2. Grayscale display of 20 consecutive frames of recorded native uint16 early endosomes from the public Ferri et al. ee01 live cell confocal acquisition, source files ee01_0001.tif through ee01_0020.tif. Frames contain CellLight Rab5a-GFP early endosomes in live HeLa cells. One first to 99.5th percentile intensity range is applied across the entire movie. The annotation is a zero-based frame index. Playback is 5 frames per second for viewing only and is independent of the acquisition timing. The MP4 is an 8-bit lossy visualization. The associated single channel TCYX TIFF is composed of the source pixels without intensity changes or a duplicated channel. All 20 frames were verified against the original TIFFs. Paper DOI: 10.1038/sdata.2018.191; Dataset DOI: 10.6084/m9.figshare.6715550.v1; Data license: CC0.

## Supplementary package installation

Install the pinned 2.5.0 wheel in a virtual environment. Use amt-refined, or import RefinedConfig and run_pipeline from automorphotrack.refined, for the particle pipeline. The amt-validated command retains the baseline implementation for Figures 2, 3, and 5-17 and Supplementary Figures 1-9. Supplementary Figure 10 reports the new particle evaluation. The automorphotrack and amt commands remain unchanged. The compressed-tiff extra supplies a TIFF decoder.

Run the examples in the source-release root or in the software/AutoMorphoTrack_release_2_5 directory from the reproduction archive. Use a virtual environment to install the pinned wheel. The particle route uses amt-refined, and the amt-validated route runs the retained baseline. Notebooks and wheels from earlier versions are kept in the archive and do not impact use of the current package.

~~~
python -m pip install “automorphotrack==2.5.0”
python -m pip install “automorphotrack[compressed-tiff]==2.5.0”
amt-refined recording.tif --out results --config configuration.json
~~~

## Particle pipeline example

The particle example uses a two-channel synthetic movie with known motion. Run the guarded adapter from the release directory after installing the 2.5.0 wheel. It checks the 18 baseline and five particle modules before importing the numerical implementation. Use a new output directory for every run. For a static image, set min_observations to 1. Particle support masks have not been validated to be mitochondrial membrane boundaries. The default setting requires three observations, thus short-lived events will not be retained and retained masks and motion summaries will not reflect these. Raw outputs preserve these events.

~~~
python -m venv .venv
# macOS and Linux:
.venv/bin/python -m pip install release_artifacts/2.5.0/automorphotrack-2.5.0-py3-none-any.whl
.venv/bin/python -I examples/refined_verified.py --input validation/refined_motion.ome.tif --out new_particle_output --config validation/refined_motion_config.json --validation-dir validation
.venv/bin/python -m pip install jupyterlab ipykernel
.venv/bin/python -m jupyter lab examples/Refined_Pipeline.ipynb
# Windows PowerShell:
.venv\Scripts\python.exe -m pip install release_artifacts/2.5.0/automorphotrack-2.5.0-py3-none-any.whl
.venv\Scripts\python.exe -I examples/refined_verified.py --input validation/refined_motion.ome.tif --out new_particle_output --config validation/refined_motion_config.json --validation-dir validation
.venv\Scripts\python.exe -m pip install jupyterlab ipykernel
.venv\Scripts\python.exe -m jupyter lab examples/Refined_Pipeline.ipynb
~~~

Minimal Debian and Ubuntu systems may need libxcb1, libgl1, and libglib2.0-0 for the OpenCV import. The GitHub CI matrix completed wheel and source installations on Windows, Linux, and macOS with Python 3.11 and 3.12, and retained test reports, dependency versions, source checks, and known-motion outputs.

~~~
sudo apt-get update
sudo apt-get install libxcb1 libgl1 libglib2.0-0
~~~

Before analyzing new data, run the validation/platform_check.py script with the validation directory provided. It validates the source modules and compares 19 baseline and 19 particle outputs with fixtures. Integer masks must be an exact match. Numerical tolerances are 10^-12 for the baseline and 10^-9 for the particle route. Retain the validation reports and use a new output directory. These checks ensure numerical reproducibility of the fixtures, not the new biological data.

~~~
# macOS and Linux:
.venv/bin/python -I validation/platform_check.py --validation-dir validation --output new_validation_check
# Windows PowerShell:
.venv\Scripts\python.exe -I validation/platform_check.py --validation-dir validation --output new_validation_check
~~~

When inputting new data, include verification of spatial or temporal calibration in the configuration file. Verify that the axes and channel assignments are correct. Inspect masks, labels, and ambiguity flags, and verify the run manifest prior to interpreting the measurements. The tested wheel and integration helper should not be modified when asking the LLM to construct a pipeline. Allow the assistant to write an external adapter around the documented functions. Compare the output of the adapter with a direct call on a known fixture. Preprocessing, parameters, datasets, and downstream statistics need to be scientifically assessed, even if the package bytes match.

**Supplementary Table 5.** Recorded integration checks.

| **Route** | **Recorded output** | **Evidence file** |
|---|---|---|
| Gemini-generated adapter, locally executed | 19 CSV/TIFF outputs exact versus standalone; maximum difference 0; speed 1 px/frame, both channels | integration/gemini_app_test/verification.json |
| Codex-operated direct and guarded Python | 19 fixture files and 20 Sample8 files exact between routes; retained Sample8 difference 8.88e-16; eight rejection controls | analysis/integration_verification.json |
| Codex-operated notebook | All three code cells passed; speed 1 px/frame, both channels | analysis/notebook_smoke_result.json |
| Expanded installation fixture | Direct and guarded outputs exact across 19 files; speed 1 px/frame, both channels | analysis/inline_revision_20261113/installation_fixture/fixture_verification.json |
| Gemini app chart renderer, Codex local execution | 23 figures; two successful runs; plotted data exact; repeated PNG/TIFF pixels exact; font and compatibility warnings retained | analysis/gemini_plot_reproduction_20261113/verification_final.json; task_status.json |
| 2.5.0 Gemini adapter and Codex CLI/guard | 19 particle output files exact; corrected metadata error; invalid manifest and changed-source controls rejected | analysis/refinement_20261114/integration/completion.json |
| 2.5.0 executed Jupyter notebook | Three code cells passed; 19 particle output files exact | analysis/refinement_20261114/integration/notebook_completion.json |
| 2.5.0 Windows, Linux, macOS CI | 12 wheel/source installations; 72 tests each; 23 modules and 38 data outputs checked | analysis/refinement_20261114/github_jobs_passed.json; GitHub run 43602083124 |
| Gemini particle figure, Codex checks | Raw indentation failure retained; repaired figure preserves 18 paired movie values and 289 accepted trace segments | outputs/refinement_20261114/figure_validation.json |

**Supplementary Table 6.** Biological tracking by movie.

| **Movie** | **Manual tracks** | **Default localization recall** | **Default accepted-link recall** | **Top-hat localization recall** | **Top-hat accepted-link recall** |
|---|---|---|---|---|---|
| MSP01 | 45 | 0.101 | 0.060 | 0.058 | 0.047 |
| MSP02 | 17 | 0.232 | 0.072 | 0.569 | 0.433 |
| MSP03 | 15 | 0.274 | 0.094 | 0.263 | 0.138 |
| MSP04 | 26 | 0.000 | 0.000 | 0.279 | 0.225 |
| MSP05 | 21 | 0.105 | 0.068 | 0.271 | 0.211 |
| MSP06 | 18 | 0.000 | 0.000 | 0.321 | 0.207 |
| MSP07 | 13 | 0.000 | 0.000 | 0.027 | 0.017 |
| MSP08 | 43 | 0.000 | 0.000 | 0.115 | 0.072 |
| MSP09 | 6 | 0.000 | 0.000 | 0.329 | 0.221 |
| MSP10 | 29 | 0.093 | 0.019 | 0.204 | 0.074 |
| MSP11 | 35 | 0.000 | 0.000 | 0.394 | 0.301 |
| MSP12 | 16 | 0.000 | 0.000 | 0.441 | 0.257 |

**Supplementary Table 7.** Mean synthetic fragmentation and coverage.

| **Configuration** | **Fragments per sequence** | **Accepted fragments per sequence** | **Missed tracks per sequence** | **Localization recall** | **Accepted-link recall** |
|---|---|---|---|---|---|
| Historical IDs | 21.80 | 21.80 | 1.52 | 0.449 | 0.156 |
| Reference masks, 6 px gate | 0.00 | 3.84 | 0.00 | 1.000 | 0.985 |
| Revised default | 3.76 | 33.72 | 1.16 | 0.575 | 0.178 |
| Threshold factor 0.6 | 1.70 | 2.88 | 1.82 | 0.412 | 0.318 |
| Threshold factor 1.0 | 2.08 | 4.38 | 0.98 | 0.724 | 0.636 |
| 6 px gate | 1.96 | 3.54 | 1.16 | 0.575 | 0.483 |
| Local 21 px | 1.66 | 2.96 | 0.00 | 0.921 | 0.870 |
| Minimum area 8 px | 1.90 | 2.50 | 1.16 | 0.575 | 0.508 |
| No opening | 1.04 | 1.74 | 2.32 | 0.361 | 0.307 |
| Top-hat 12 px | 1.98 | 4.28 | 0.34 | 0.850 | 0.766 |

**Supplementary Table 8.** Sample8 runtime and memory.

| **PNG rendering** | **Processes** | **Median time (s)** | **Time range (s)** | **Median peak RSS (MiB)** |
|---|---|---|---|---|
| Disabled | 3 | 2.21 | 2.21 to 3.02 | 336.4 |
| Enabled | 3 | 2.42 | 2.42 to 2.67 | 410.3 |

**Supplementary Table 9.** Reserved-movie particle tracking at 3 pixels.

| **Movie** | **Manual tracks** | **Top-hat link recall** | **Particle localization recall** | **Particle link recall** | **Particle missed / fully recovered** |
|---|---|---|---|---|---|
| MSP04 | 26 | 0.225 | 0.569 | 0.500 | 3 / 2 |
| MSP05 | 21 | 0.211 | 0.821 | 0.693 | 1 / 0 |
| MSP06 | 18 | 0.207 | 0.888 | 0.790 | 2 / 3 |
| MSP10 | 29 | 0.074 | 0.491 | 0.408 | 7 / 3 |
| MSP11 | 35 | 0.301 | 0.940 | 0.911 | 0 / 17 |
| MSP12 | 16 | 0.257 | 0.901 | 0.875 | 0 / 8 |

