## Supplementary material for "AutoMorphoTrack: A modular framework for quantitative analysis of organelle morphology, tracking, and image colocalization": Validated Core and Visual Guide

#### Current core and integration

When requesting the integration of the particle pipeline, use the 2.5.0 example and keep the numerical package and both source manifests unchanged. Request an external adapter, explicit configuration, runnable commands, and a comparison with the supplied fixture. Record other processes like preprocessing and post-pipeline calculations. Source checks do not validate choices for preprocessing or post-pipeline calculations. The example prompt below describes the current route.

#### Figure 4 Integration architecture

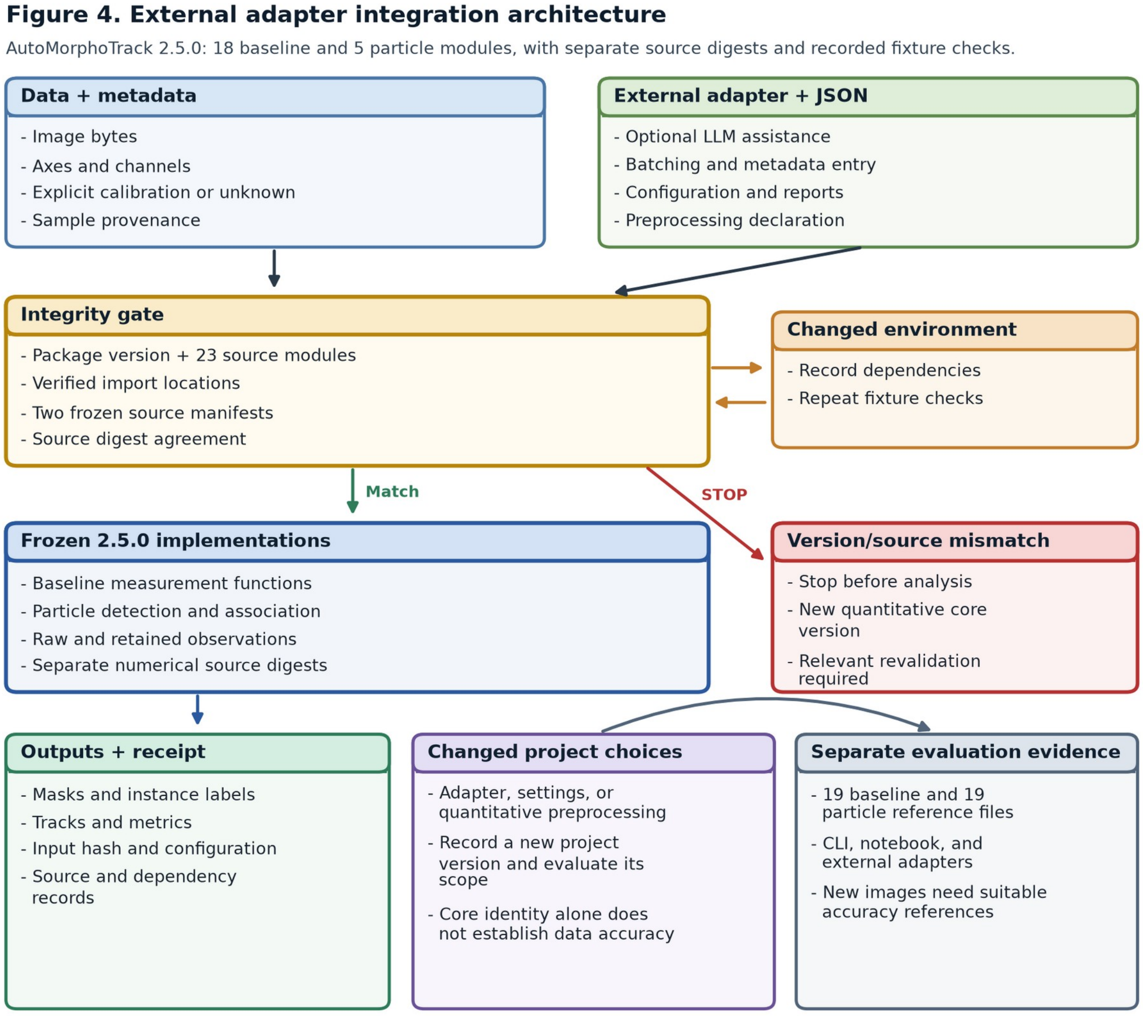


The input data and metadata are routed to the 2.5.0 particle pipeline via an external adapter using JSON configuration. An example helper checks the installed package version, both source manifests covering 23 modules, and import locations. A source mismatch stops the analysis. The run manifest captures input identity, settings, source and dependency versions, observation counts, and output checksums. New adapters, settings, preprocessing, and environments require new checks. The tested execution paths are supported by the supplied reference outputs; however, they do not validate output correctness for new data.

### Figure 2 Public confocal workflow

Figure 2. Public confocal image workflow

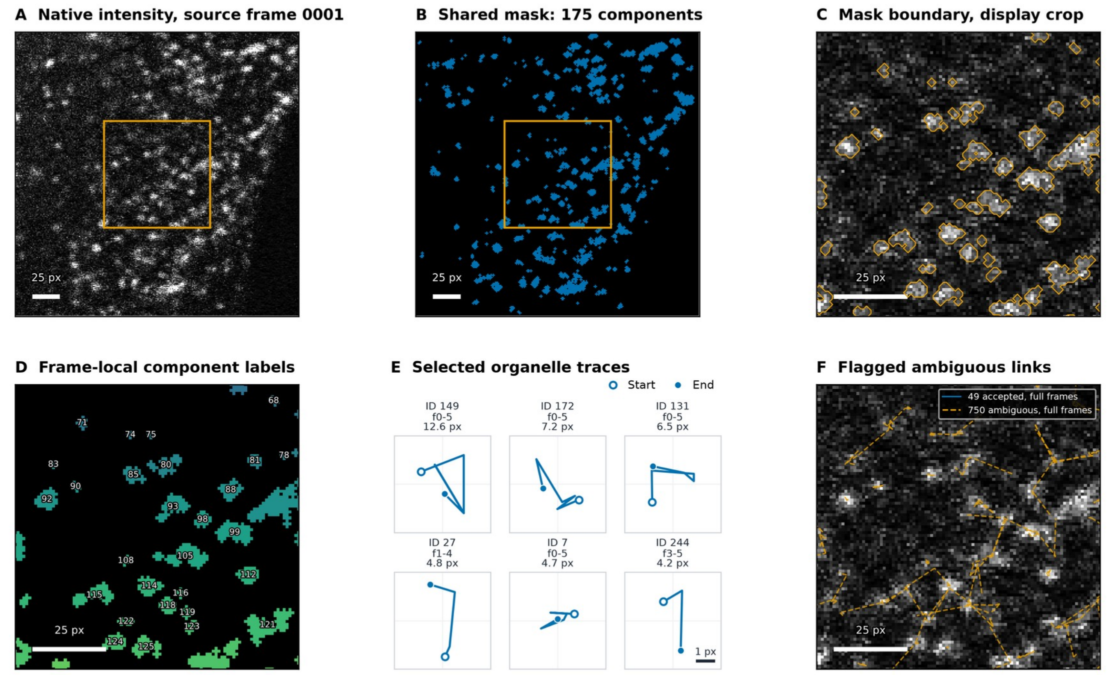


Ferri et al. 2018; early endosomes, live HeLa cells; acquisition ee01; CC0.
A-B: full field; C, D, F: central display crop. E: six longest continuous accepted segments, each at least two steps.
E-F: source frames 0001-0006; 15 px gate. Trace labels use zero-based frames. Ambiguous links are excluded.

Public early-endosome confocal images from Ferri et al. (2018), acquisition ee01, demonstrate the revision functions on native uint16 data. A shows source frame ee01_0001.tif. B shows the foreground mask associated with ee01_0001.tif and contains 175 full frame components. Orange boxes show the central display crop (80-175 rows and 80-175 columns; zero-based indices) shown in C, D, and F. C shows the foreground mask boundaries and D labels connected components, with labels local to that frame and omitted if text would cross the edge of the display. E displays the six continuous accepted trace segments with the longest path length in source frames 0001 to 0006, each with at least two accepted steps. Traces are centered in 5-pixel-wide views at the same scale. Rotation and smoothing have not been applied. Markers identify the beginning and end of each trace; labels provide track ID, zero-based frame range, and path length. Ambiguous links and gaps end a trace segment. These examples have been chosen to illustrate movement. F shows accepted links in blue and flagged ambiguous assignments in orange dashed lines over source frame 0001. Across the full six-frame fields there are 49 accepted links and 750 flagged ambiguous assigned links; crop boundaries clip their display. The default segmentation uses Otsu factor 0.8, a one-pixel opening disk, removal of components under three pixels, border clearing and eight-connected component labeling. The link gate is set to 15 pixels. These examples show the execution and uncertainty flags. No independent masks or trajectories are provided. Images are displayed using the same frame-0001 first-to-99.5th percentile limits and do not change measurement intensity. Image scale bars are 25 pixels; trace views in E are 5 pixels wide. Data DOI: 10.6084/m9.figshare.6715550.v1; paper DOI: 10.1038/sdata.2018.191; CC0.

### Figure 3 Controlled translation quality control

**Figure 3. Controlled translation QC on a public image**

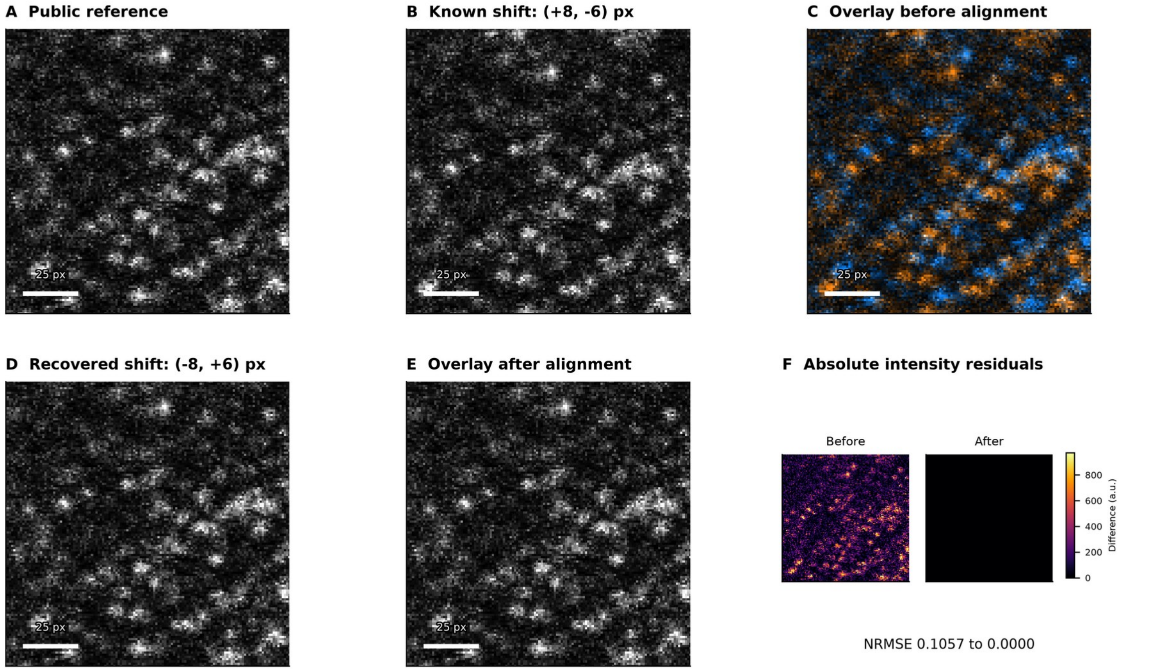


The moving image is a known transform of ee01_0001.tif, not a second native acquisition frame.
Reference is blue and moving is orange in overlays; agreement is neutral. Panels show the same central 128 x 128 px crop.
QC uses 58,560 common valid pixels; bilinear resampling applies only to the second image. Primary measurements are unchanged.

##### Supplementary Movie 2

Supplementary Movie 2. Grayscale display of 20 consecutive frames of recorded native uint16 early endosomes from the public Ferri et al. ee01 live cell confocal acquisition, source files ee01_0001.tif through ee01_0020.tif. Frames contain CellLight Rab5a-GFP early endosomes in live HeLa cells. One first to 99.5th percentile intensity range is applied across the entire movie. The annotation is a zero-based frame index. Playback is 5 frames per second for viewing only and is independent of the acquisition timing. The MP4 is an 8-bit lossy visualization. The associated single channel TCYX TIFF is composed of the source pixels without intensity changes or a duplicated channel. All 20 frames were verified against the original TIFFs. Paper DOI: 10.1038/sdata.2018.191; Dataset DOI: 10.6084/m9.figshare.6715550.v1; Data license: CC0.

#### Protected technical appendix

#### Example prompt for LLM-assisted integration

```
Write a project-specific Python adapter and JSON configuration around AutoMorphoTrack 2.5.0. Use
examples/refined_verified.py::run_verified in a fresh isolated Python process. Keep the installed
automorphotrack.validated and automorphotrack.refined modules, the helper,
validation/core_manifest.json, and validation/refined_manifest.json unchanged.

Organize input files, collect explicitly supplied metadata, batch analyses, and produce reports
from saved outputs. Do not invent calibration, channel identity, biological replication, or
ground truth. Require explicit axes and channel assignments. Record preprocessing, settings,
dependencies, and downstream calculations separately. Source integrity does not establish
biological accuracy.

Run the supplied validation/refined_motion.ome.tif fixture using
validation/refined_motion_config.json before analyzing project data. Compare the adapter outputs
with validation/refined_motion_expected, excluding the timestamp-bearing run_manifest.json from
byte comparisons. Compare numerical CSV values and integer TIFF arrays, and retain all failures.
Use a new output directory for each run. Verify 19 particle data files and preserve raw as well
as retained tracks and labels.

Supply the adapter source, configuration, preprocessing record, commands, results, and
limitations. Do not edit or replace the packaged numerical functions. Changes to those functions
require a new version and relevant revalidation. Do not add a chatbot to the analysis runtime or
upload microscopy data to an external service.
```

#### Runnable command examples

```
.venv/bin/python -I examples/refined_verified.py --input validation/refined_motion.ome.tif --out
new_adapter_check --config validation/refined_motion_config.json --validation-dir validation
```
