## Supplemental Tables 1-4 for "AutoMorphoTrack: A modular framework for quantitative analysis of organelle morphology, tracking, and image colocalization"

### AutoMorphoTrack Supplementary Tables 1 to 4

#### Supplementary Table 1 Software scope and implementation

The measured accuracy comparison is between the legacy and revised AutoMorphoTrack implementations. Head-to-head accuracy against the named tools was not tested.

| Software | Primary scope | Interface or implementation | Relation to this revision | Source |
|---|---|---|---|---|
| AutoMorphoTrack 2.5.0 baseline | 2D two-channel morphometry, centroid linking, colocalization | automorphotrack.validated; amt-validated; original 1.1.0rc2 core | 18 unchanged source modules; shared labels; gated one-to-one links; recorded configuration | Manuscript reference 22; GitHub release v2.5.0 |
| AutoMorphoTrack 2.5.0 particle | 2D particle detection and tracking; one or more channels; colocalization requires two | automorphotrack.refined; amt-refined; five particle source modules | Gaussian-background removal; multiscale LoG detection; distance, intensity and area association costs | Manuscript reference 22; Supplementary Figure 10 and Table 9 |
| CellProfiler | Modular image and object analysis, tracking, colocalization | Open-source graphical pipeline environment | Broader configurable pipeline platform | https://doi.org/10.1186/s12859-021-04344-9 |
| Imaris | 3D and 4D visualization, segmentation, tracking | Commercial graphical analysis platform | Multidimensional analysis platform | https://imaris.oxinst.com/products/imaris-for-cell-biologists |
| MiNA | Mitochondrial footprint and skeleton morphology | Fiji workflow | Network morphology and branch measurements | https://doi.org/10.1016/j.acthis.2017.03.001 |
| MitoGraph | 3D mitochondrial volume and network topology | Open-source image-analysis software | Three-dimensional network quantification | https://doi.org/10.1016/bs.mcb.2014.10.003 |
| Mitochondria Analyzer | Confocal mitochondrial morphology and function | Fiji plugin | Mitochondrial preprocessing and measurement | https://doi.org/10.1152/ajpendo.00457.2019 |
| Cellpose | Learned cellular instance segmentation | Open-source learned model | External labels can be passed to the AutoMorphoTrack baseline route | https://doi.org/10.1038/s41592-020-01018-x |
| StarDist | Learned star-convex instance segmentation | Open-source 2D and 3D models | External labels can be passed to the AutoMorphoTrack baseline route | https://github.com/stardist/stardist |

#### Supplementary Table 2 Public datasets

| Study and article | Selected inputs | Reference and scope | Data source and license |
|---|---|---|---|
| Ferri 2018<br>Article DOI:<br>10.1038/sdata.2018.191 | ly01, ly02, ee01, ee02, le01, le02; first 20 frames of each acquisition | None; execution audit only | https://doi.org/10.6084/m9.figshare.c.4070033<br>CC0 1.0 |
| MitoLandscape 2025<br>Article DOI:<br>10.3389/fcell.2025.1668779 | example_MITO.tif; static maximum-intensity projection of 30 Z planes | Source MITOBIN outputs archived but not treated as ground truth | https://zenodo.org/records/16037033<br>CC BY 4.0 |
| OrganelleProfiler 2023<br>Article DOI:<br>10.1371/journal.pone.0278009 | Two 4-channel OCP fields (VWF with PDI or Rab27A) and one EEA1 RGB export; five 2D channel representations | None; source CellProfiler results are not independent masks | https://zenodo.org/records/7875537<br>CC BY 4.0 |
| mtFociCounter 2023<br>Article DOI:<br>10.1093/nar/gkad864 | First four raw TIFF members; static channel 1 Z projections | None; manual cell segmentation does not provide mitochondrial masks | https://zenodo.org/records/7634536<br>CC BY 4.0 |

### Supplementary Table 2 Public datasets continued

| Study and article | Selected inputs | Reference and scope | Data source and license |
|---|---|---|---|
| MoDL 2025<br>Article DOI: 10.1038/s41467-025-55825-x | 0000 through 0009 image-mask pairs; 0003 through 0009 held out from AMT development; seven held-out patches used for the reported independent mask evaluation | Published binary semantic training masks; paper-level manual annotation provenance; not independent expert adjudication | https://github.com/OBPNPW2024/MoDL/tree/1053e473785560bc473aa64fd357379f9f0e21e2/deform<br>GPL-3.0 repository; no separate per-image license is stated<br>The linked Zenodo application distribution is CC BY 4.0, but matched source training pairs were not located there. That record is not used to relabel these GitHub images as CC BY. |
| Myofibril 2021<br>Article DOI: 10.1038/s41467-021-22058-7 | All 13 T0 image-mask pairs; native 128x128 patches | Author-drawn Fiji masks; one parent stack; source-model training data | https://github.com/fabda/Myofibril_paper/tree/11f512a8181fb98540b6e5a89466ed85b46b4d5f<br>GPL-3.0 repository; separate linked image archive license unspecified<br>Raw linked archive excluded from redistributable package; acquisition URL and hashes retained |
| MitoHacker 2020<br>Article DOI: 10.1038/s41598-020-75899-5 | All 17 original multicell TIFFs; green channel of RGB exports | None; execution and parameter sensitivity only | https://github.com/Mitogenie/Mito-Hacker/tree/18765f1d584db04be77f3397f730b98a51225394<br>MIT repository; no separate per-image license stated |
| MSP tracker 2025<br>Article DOI: 10.1371/journal.pbio.3003099 | Version 1; 12 deposited Airyscan and spinning-disk movies; duplicate Fas3 export excluded; six development and six reserved movies for the particle route | 284 distinct deposited trajectories; paper reports 391. Partial manual annotations; unmatched predictions remain unverified. Six reserved movies contain 145 trajectories and 2,764 adjacent reference links. | https://doi.org/10.17863/CAM.114338<br>CC BY 4.0<br>Reserved movies were previously evaluated with the baseline; within-study evaluation. |

### Supplementary Table 3 Worked example observations

Baseline route in AutoMorphoTrack 2.5.0; 40 frames per region from recording no_PFF_Cont003. Repeated detections are frame-object observations. The regions share one recording.

| Region | Organelle | Detections | Candidate track IDs | Tracks with accepted steps | Accepted links | Ambiguous links |
|---|---|---|---|---|---|---|
| Sample1 | Mitochondria | 2359 | 508 | 108 | 498 | 1353 |
| Sample1 | Lysosomes | 2068 | 276 | 81 | 518 | 1274 |
| Sample2 | Mitochondria | 1785 | 391 | 117 | 530 | 864 |
| Sample2 | Lysosomes | 2525 | 348 | 78 | 471 | 1706 |
| Sample3 | Mitochondria | 1961 | 436 | 84 | 465 | 1060 |
| Sample3 | Lysosomes | 2962 | 363 | 92 | 632 | 1967 |
| Sample4 | Mitochondria | 1887 | 407 | 89 | 363 | 1117 |
| Sample4 | Lysosomes | 2787 | 295 | 83 | 428 | 2064 |
| Sample5 | Mitochondria | 1736 | 326 | 85 | 468 | 942 |
| Sample5 | Lysosomes | 1600 | 228 | 74 | 452 | 920 |
| Sample6 | Mitochondria | 1800 | 423 | 94 | 410 | 967 |
| Sample6 | Lysosomes | 2030 | 277 | 74 | 446 | 1307 |
| Sample7 | Mitochondria | 1717 | 347 | 84 | 310 | 1060 |
| Sample7 | Lysosomes | 1281 | 238 | 50 | 173 | 870 |
| Sample8 | Mitochondria | 2084 | 427 | 79 | 333 | 1324 |
| Sample8 | Lysosomes | 2344 | 246 | 73 | 524 | 1574 |
| Sample9 | Mitochondria | 2127 | 410 | 98 | 500 | 1217 |
| Sample9 | Lysosomes | 1303 | 197 | 57 | 378 | 728 |
| Sample10 | Mitochondria | 1442 | 274 | 72 | 429 | 739 |
| Sample10 | Lysosomes | 1379 | 179 | 48 | 313 | 887 |

### Supplementary Table 4 Implementation changes

Baseline revisions are retained in automorphotrack.validated. The final three rows identify the separate 2.5.0 particle route and preservation of the baseline core.

| Area | Historical implementation | Revision | Evidence |
|---|---|---|---|
| Segmentation consistency | Module-specific masks and parameter choices | One shared label set per channel | Regression tests; saved masks and instance labels |
| Segmentation options | Global thresholding and cleanup | Global/local thresholding; white top-hat; external instance labels | Fixed synthetic and public-data comparisons |
| Intensity and axes | Display-derived examples; guessed channel layouts | Native numeric intensity; explicit validated axes | Native16 reconstruction; axis regression tests |
| Temporal identity | Object IDs restarted per frame | Persistent IDs; gated one-to-one assignment | Identity controls; all 50 synthetic sequences |
| Tracking uncertainty | Competing links not distinguished consistently | Ambiguity flags; accepted-step summaries | Precision/recall and coverage reported together |
| Motion quantities | Displacement and velocity definitions mixed | Path length; conditional net displacement; calibrated or px/frame units | Known 1 px/frame fixture; regression tests |
| Circularity | Mixed raster area/perimeter estimator | Matched contour area/perimeter; no clipping | 7,167/12,505 vs zero above 1 on identical public-data masks |
| Colocalization | Binary/intensity definitions and names inconsistent | Separate mask fractions, Jaccard, intensity Pearson, cosine overlap | Known-value and undefined-input controls |

### Supplementary Table 4 Implementation changes continued

| Area | Historical implementation | Revision | Evidence |
|---|---|---|---|
| Integrated summary | Duplicated frames; identifier correlations | One row per frame; Spearman; paired coverage | Regression tests; Frame_Correlation_N.csv |
| Plot support | Unbounded kernel-density tails | Empirical cumulative distributions | Regenerated figures and underlying finite values |
| Distribution | Broken or unsupported install routes | Pinned 2.5.0 wheel and source distribution; baseline and particle CLI routes | 72 tests per installation; 12 operating-system, Python and distribution combinations |
| Reproduction | Incomplete settings and source identities | Input, code, environment, settings and output receipts | Run manifests and numerical-equivalence checks |
| External LLM integration | Native chatbot capability implied without implementation | External adapters; installed-version and source-integrity checks before numerical imports | Recorded Gemini source execution; Codex and Jupyter results; 19 particle reference output files |
| Visual alignment check | No tested standalone alignment illustration | Separate phase-correlation QC helper; primary inputs unchanged | Five known transforms; ten helper tests |
| Particle detection in 2.5.0 | Global and fixed top-hat threshold comparators | Gaussian-background subtraction; scale-normalized LoG response; watershed support masks | Six reserved biological movies; 30 new synthetic movies; five noise-only controls |
| Particle association in 2.5.0 | Distance-only centroid linking | Distance-normalized assignment with intensity, area, and birth/death costs; three-observation retention | Mean accepted-link recall 0.696 versus 0.212 for fixed top-hat on six reserved movies |
| Baseline preservation in 2.5.0 | Validated 1.1.0rc2 numerical core | 18 unchanged baseline modules; five separately frozen particle modules | Two source manifests; baseline and particle fixture outputs; package 2.5.0 identity |
